# Landscape change and alien invasions drive shifts in native lady beetle communities over a century

**DOI:** 10.1101/2022.04.21.489069

**Authors:** Kayla I Perry, Christie A Bahlai, Timothy J Assal, Christopher B Riley, Katherine J Turo, Leo Taylor, James Radl, Yvan A Delgado de la flor, Frances S Sivakoff, Mary M Gardiner

## Abstract

**Aim:** Understanding drivers of insect population declines is essential for the development of successful conservation plans, but data limitations restrict assessment across spatial and temporal scales. Museum records represent a unique source of historical data that can be leveraged to investigate temporal trends in insect communities. Native lady beetle decline has been attributed to competition with established alien species and landscape change, but the relative importance of these drivers is difficult to measure with short-term field-based studies. Here we assessed distribution patterns for native lady beetle species over 12 decades using museum records and evaluated the relative importance of alien species and landscape change as long-term drivers contributing to changes in lady beetle communities.

**Location:** Ohio, USA.

**Methods:** We compiled occurrence records for 28 lady beetle species collected in Ohio, USA from 1900-2018. Incidence-based measures of taxonomic beta-diversity were used to evaluate changes in lady beetle community composition over time. To evaluate the relative influence of temporal, spatial, landscape, and community drivers on the captures of native lady beetles, we constructed negative binomial generalized additive models.

**Results:** We report evidence of declines in captures for several native species, including *Adalia bipunctata, Coccinella novemnotata, Hippodamia convergens*, and *Coleomegilla maculata*. Importantly, the timing, severity, and drivers of these documented declines were species-specific. Changes in lady beetle species composition began in the 1980s, when processes of species loss/gain and turnover shifted communities towards dominance by a few alien lady beetle species. Land cover change also was associated with declines in captures, particularly for *C. novemnotata* which declined prior to the arrival of alien species in the state.

**Main conclusions:** Our study documented shifts in Ohio’s lady beetle communities beginning in the 1980s as alien species supplanted natives. Drivers of declines in captures of native lady beetle species were highly species-specific, emphasizing that mechanisms driving population losses cannot be generalized even among closely related species. These findings also indicate the importance of museum holdings and the analysis of species-level data when studying temporal trends in insect populations.

## Introduction

Global biodiversity loss is a growing threat to the ecosystem function and services for which humans depend (Dirzo & Raven, 2003; Mace *et al*., 2012; Ceballos *et al*., 2015; Cardoso *et al*., 2020). Given the importance of insects for ecosystem services such as pollination, pest suppression, and nutrient cycling (Losey & Vaughan, 2006; Yang & Gratton, 2014), understanding the causes of documented spatiotemporal changes in insect populations is a critical research focus. Numerous recent studies have pointed to declines in the abundance, richness, and biomass of insects (Hallmann *et al*., 2017; Seibold *et al*., 2019), including bees (Grixti *et al*., 2009; Potts *et al*., 2010), beetles (Brooks *et al*., 2012; Homburg *et al*., 2019), leafhoppers and planthoppers (Schuch *et al*., 2012), and butterflies and moths (Maes & Van Dyck, 2001; Conrad *et al*., 2006; Habel *et al*., 2016; Warren *et al*., 2021). Although population declines have been frequently reported, relatively stable or increasing populations of insects also have been documented in some settings (Shortall *et al*., 2009; Fox *et al*., 2014; Crossley *et al*., 2020; Schowalter *et al*., 2021). For example, observations of moths in Great Britain identified highly species-specific temporal patterns over a 40 year period, as 260 species declined in frequency, 160 species increased, and 253 species remained unchanged (Fox *et al*., 2014). Complexity in the responses of insects has generated scientific debate about whether observed declines are generalizable across insect taxa and guilds and at larger spatial scales (Simmons *et al*., 2019; Thomas *et al*., 2019; Didham *et al*., 2020; Saunders *et al*., 2020). Although many challenges to studying insect population trends exist (Didham *et al*., 2020; Montgomery *et al*., 2020), understanding the magnitude and drivers of declines in insect species is critical for biodiversity conservation management and the maintenance of insect-based ecosystem services and function.

One primary limitation to understanding insect populations over time stem from data deficiencies such as low taxonomic resolution, geographic restrictions, and short time series (Sánchez-Bayo & Wyckhuys, 2019; Thomas *et al*., 2019; Didham *et al*., 2020). However, specimen records from natural history museums and other institutions can be leveraged to investigate trends in insect populations over greater spatial and temporal scales (Lister, 2011; Kharouba *et al*., 2019). Although specimen record-based data have their own set of biases and challenges (Boakes *et al*., 2010; Meineke & Daru, 2021), biological collections represent a unique source of historical data that documents the occurrence of species in time and space. For example, museum records revealed that 11 bumble bee species native to eastern North America and Canada have experienced substantial (>50%) declines in captures over the past century, while eight species have remained stable or increased in collections (Colla *et al*., 2012). Importantly, specimen records can be used as baseline measures for investigating the impacts of anthropogenic drivers such as the establishment of exotic species, environmental degradation, and climate change on patterns of biodiversity (Suarez & Tsutsui, 2004; Winker, 2004; Johnson *et al*., 2011; Kharouba *et al*., 2019). For instance, moth assemblages collected on Mount Kinabalu in Borneo in 2007 were compared to historical records collected from the same sites in 1965, revealing upward shifts in the altitudinal distribution of species in response to changes in temperature (Chen *et al*., 2009). Although the use of collections data is limited in its ability to track the absolute abundance of a species due to sampling effort variation (Ries *et al*., 2019), historical specimen data facilitate understanding of species’ responses to anthropogenic change by helping to distinguish signals of decline from natural population variability, especially when considering relative observations in groups of similar taxa. Therefore, specimen records are invaluable resources that can be harnessed to address biodiversity conservation initiatives, including documenting changes in communities of beneficial insects.

Lady beetles (Coleoptera: Coccinellidae) are a family of charismatic insect species that are commonly collected and contribute broadly to pest suppression by consuming aphids, scales, psyllids, mites, fungi, and other pests (Evans, 2009; Hodek & Honěk, 2009; Weber & Lundgren, 2009; Hodek *et al*., 2012). Because lady beetles are capable of rapidly colonizing habitats to exploit ephemeral prey resources, these species have been used widely in biological control programs in agricultural systems (Caltagirone & Doutt, 1989; Obrycki & Kring, 1998; Koch, 2003; Rondoni *et al*., 2021). A number of authors have noted declines in native lady beetle populations across the United States (Alyokhin & Sewell, 2004; Harmon *et al*., 2007) and Europe (Roy *et al*., 2012; Brown & Roy, 2018), which may contribute to an overall loss of resilience of the biological control services offered by this community (Bahlai *et al*., 2021). For example, the historically widespread native aphidophagous species *Hippodamia convergens* Guerin has declined in the US states of Michigan (Gardiner *et al*., 2009), Ohio (Gardiner *et al*., 2012), Wisconsin (Steffens and Lumen 2015), and Minnesota (Steffens and Lumen 2015), as well as the Canadian Province of Manitoba (Turnock et al. 2003). Likewise, the nine-spotted lady beetle *Coccinella novemnotata* Herbst was once common in eastern North America but had not been collected for over a decade until a community scientist “rediscovered” it (Losey *et al*., 2007). Anthropogenic activities such as the establishment of alien species and landscape change have been hypothesized as potential drivers of native lady beetle decline (Alyokhin & Sewell, 2004; Harmon *et al*., 2007; Gardiner *et al*., 2012; Honek *et al*., 2014; Bahlai *et al*., 2015; Roy *et al*., 2016).

Several studies have observed that the decline of native aphidophagous species coincided with the establishment and spread of alien lady beetle species, particularly the Asian species *Harmonia axyridis* (Pallas) and the European species *Coccinella septempunctata* (Linnaeus) (Turnock *et al*., 2003; Roy *et al*., 2012; Steffens & Lumen, 2015; Roy *et al*., 2016). Following their establishment, alien lady beetles became the dominant species within many native communities (Alyokhin & Sewell, 2004; Harmon *et al*., 2007; Bahlai *et al*., 2015). Because of their dominance, direct and indirect competitive interactions with alien species are hypothesized as a driver of declines in native lady beetles (Pell *et al*., 2008; Li *et al*., 2021). For example, intraguild predation has been documented among native and alien lady beetles in the field (Gagnon *et al*., 2011; Thomas *et al*., 2013; Brown *et al*., 2015; Ortiz-Martínez *et al*., 2020), wherein native eggs and larvae were more likely to be the intraguild prey for alien species (Snyder *et al*., 2004; Katsanis *et al*., 2013). Apparent competition also has been observed in the field, as native species experienced greater egg predation from a guild of shared predators than alien lady beetle species (Smith & Gardiner, 2013). These asymmetric interactions may largely benefit alien species at the expense of native lady beetle populations, but the extent and context of these effects on native species are difficult to quantify. The short time scales of many studies and the lack of data from lady beetle communities before the establishment of alien species limits understanding of the impacts of these invaders.

Landscape change that results in the loss, fragmentation, and degradation of natural habitat also has been hypothesized as a key driver contributing to population declines of insect species (Potts, *et al*. 2010; Wagner, 2020; Wagner *et al*., 2021), including lady beetles (Honek *et al*., 2017). Land cover change resulting from increased urbanization and agricultural intensification can influence the structure and composition of lady beetle communities (Gardiner *et al*., 2009; Woltz & Landis, 2014; Grez *et al*., 2019; Parker *et al*., 2020). For example, native and alien species were less abundant in isolated urban greenspaces that were embedded in landscapes dominated by impervious surfaces and built infrastructure (Parker *et al*., 2020). In agricultural landscapes, native and alien lady beetles were more abundant in fields surrounded by higher crop diversity and more semi-natural habitat types such as grasslands and forests (Woltz & Landis, 2014). Loss and fragmentation of natural habitat in landscapes such as those dominated by urban and agricultural land cover may differentially affect species depending on their life history traits such as phenology, dispersal ability, overwintering biology, and food requirements (Zaviezo *et al*., 2006). However, the impacts of landscape change on native lady beetle populations may occur gradually over time, making species responses difficult to detect over short time scales. For example, gradual directional change in the species composition of native lady beetle communities was documented over a 118-year period in Missouri, USA using museum specimen records (Diepenbrock *et al*., 2016). Because there was no evidence that the establishment of alien lady beetle species affected the rate of change, Diepenbrock et al. (2016) hypothesized that these long-term community changes may be related to altered land use patterns. The substantial year to year variation in the abundance of lady beetle species (Elliott *et al*., 1996; Honek *et al*., 2014) requires longer time series data to detect changes in populations caused by landscape change and to distinguish these effects from other anthropogenic stressors (Bahlai & Zipkin, 2020; Bahlai *et al*., 2021).

While several hypotheses have been proposed, the causes of declines in native lady beetle populations remain under debate. Importantly, these hypotheses are not mutually exclusive, and it is likely that causes of declines in native species are multi-faceted wherein multiple mechanisms are responsible for the observed patterns in lady beetle populations. Although causes of declines in native species are often studied independently, these drivers may interact to influence native populations (Didham *et al*., 2007). For example, habitat modification that transitions natural habitat to more highly disturbed urban and agricultural habitats may differentially benefit alien species over native lady beetles (Grez *et al*., 2013), with implications for direct and indirect competitive interactions among native and alien species. In contrast, landscapes with less disturbed perennial grasslands can serve as an important refuge primarily for native lady beetle species, providing prey and habitat requirements necessary for survival and reproduction (Evans, 2004; Diepenbrock & Finke, 2013). Understanding the magnitude and drivers of declines in native lady beetle populations will require comprehensive time series data documenting community responses that can then be used to assess the contribution of the establishment of alien species and landscape change simultaneously.

To understand the relative importance of the establishment of alien species and landscape change as drivers of native species decline, we compiled historical occurrence records of lady beetles collected in Ohio, USA from museums and other institutions across the United States. Our goals were to assess long-term patterns in native lady beetle species occurrence and communities within the region, and to evaluate the relative importance of the establishment and spread of alien lady beetle species and landscape change as drivers contributing to native species decline in this community.

## Methods

### Lady beetle specimens and data requests

To investigate long-term changes in native lady beetle communities within Ohio, USA (Fig. 1), we used historic occurrence records for native and alien species within the tribe Coccinellini and four additional non-Coccinellini species. Targeted Coccinellini genera were *Adalia, Anatis, Anisosticta, Aphidecta, Calvia, Ceratomegilla, Coccinella, Coelophora, Coleomegilla, Cycloneda, Harmonia, Hippodamia, Macronaemia, Mulsantina, Myzia, Naemia, Neoharmonia, Olla*, and *Propylea*, and non-Coccinellini species were *Brachiacantha ursina* (Fabricius), *Chilocorus stigma* (Say), *Hyperaspis undulata* (Say), and *Psyllobora vigintimaculata* (Say). We contacted 59 institutions based within the United States, the majority of which are hosted by the Entomological Collections Network (ENC, 2020). Ohio lady beetle records were compiled from 25 institutions with assistance of their curators (see Appendix 1: Table S1.1 in Supporting Information).

**Figure 1.**
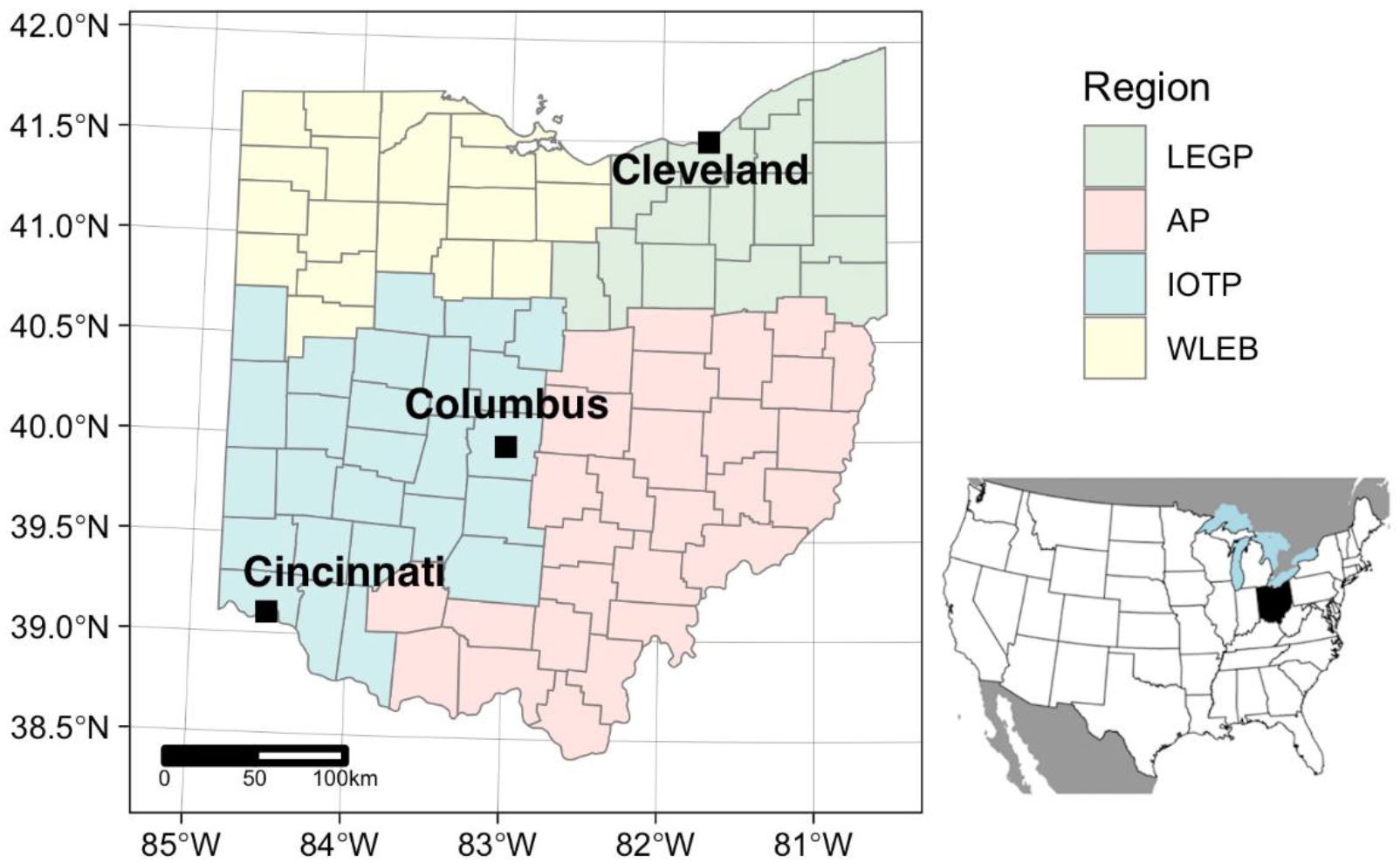
The study area is congruent with the state of Ohio; inset shows the relative location of Ohio in the conterminous United States. Ohio counties within the four geographic regions of the state are color-coded, Lake Erie Glaciated Plateau (LEGP; top right), Appalachian Plateau (AP; bottom right), Indiana-Ohio Till Plain (IOTP; bottom left), and Western Lake Erie Basin (WLEB; top left).

Specimen loans were requested from museums and institutions that had unidentified lady beetle records from Ohio. Any unidentified lady beetle species that were unaccounted for within the collections of these institutions were not included within our dataset. Lady beetle species identifications were determined using Gordon (1985). All lady beetle species were characterized by their status (native or alien to North America) and their primary diet (aphidophagous, coccidophagous, or fungivorous) which was based on most frequently reported prey (aphids, scales, or fungi) (Angalet *et al*., 1979; Gordon, 1985; Dixon & Dixon, 2000; Michaud, 2001; Staines, 2008; Hodek *et al*., 2012; Majerus, 2016).

### Land use and land cover change analysis

To assess the influence of landscape change on lady beetle communities, we analyzed historical land use and land cover (LULC) data from four points in time (1938, 1970, 1992, 2016). Annual historical LULC data were obtained from the US Geological Survey (Sohl *et al*., 2018) for the years 1938, 1970, and 1992. These historical LULC backcasts were modeled using numerous historical data sets and created explicitly to extend the National Land Cover Database (NLCD) to earlier time periods, prior to the availability of remote-sensing data (Sohl *et al*., 2016). The NLCD and historical backcast data were crosswalked into four primary land use classes: agriculture, developed, forest, and non-target (i.e., all other land cover classes found in Ohio) (see Appendix 2: Table S2.1, Table S2.2 in Supporting Information). The contemporary landscape of Ohio was assessed with the 2016 NLCD (Dewitz, 2019; Wickham *et al*., 2021). To assess change over time, we calculated the total area of each cover class, along with the percentage of area occupied by each class, for each county at each of the four time periods. Although the LULC data have different spatial resolutions (NLCD = 30 m; historical backcasts = 250 m) they were not resampled, as the derived metrics (e.g. total area of cover class, percentage of county area) are relative, and therefore comparable across time. Analyses were completed in R (R Core Team, 2020) using the ‘raster’ and ‘sf’ R packages (Pebesma, 2018; Hijmans, 2020).

There was an increase in forest cover from 1938 to 1970, and it has held steady since the early 1990s (see Appendix 2: Fig. S2.6). Most of the increase occurred in the Appalachian Plateau and Lake Erie Glaciated Plateau regions in the eastern and southern portions of the state (Fig. 2). There was a steady increase in developed land around existing population centers, with a larger increase in suburban areas since the early 1990s. The increase in forest and developed lands came at the expense of agricultural lands, yet this cover class remains the dominant land cover in many counties of the state (Fig. 2; see Appendix 2: Fig. S2.6). During each time period, there was high variability in the amount of agricultural and forested land across Ohio counties (see Appendix 2: Fig. S2.6).

**Figure 2.**
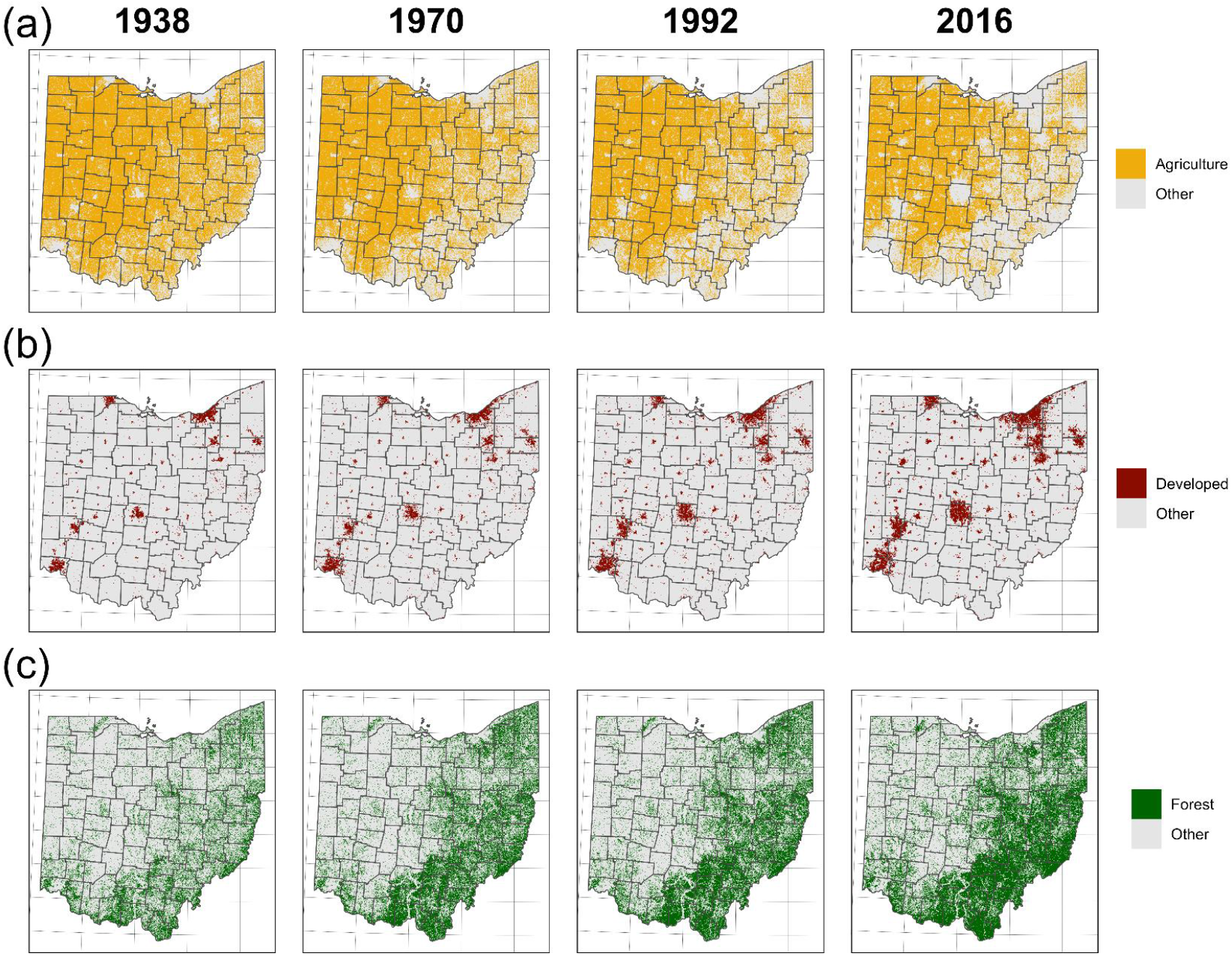
Spatial extent of agriculture (row A), developed land (row B), and forest (row C) in Ohio for each time period (columns).

Land cover data were compiled at the county and decade resolution to align with the spatial and temporal resolution available for the museum specimen data. For each county, we used an assumed linear progression between available timepoints to estimate the percentage land cover of a given class in each decade. The 1930 land cover data were extrapolated using the same procedure with available data from 1938-1970. Because lady beetle data were generally recorded at the county level, we computed the geographic centroids of each Ohio county based on present-day county boundaries as defined by the Ohio Department of Transportation (http://ogrip-eohio.opendata.arcgis.com/datasets/odot-county-boundaries).

### Statistical analysis

Lady beetle museum records were included in the analyses if they were collected within Ohio, and we could obtain county-level location data and the year of collection. All analyses were conducted in R (R Core Team, 2020).

Changes in total lady beetle and aphidophagous lady beetle community composition over time were assessed using incidence-based measures of taxonomic beta-diversity. Due to low county-level counts, county records of lady beetle species were pooled by Ohio geographic region (Fig. 1; Lake Erie Glaciated Plateau (LEGP) in the northeast, Appalachian Plateau (AP) in the southeast, Indiana-Ohio Till Plain (IOTP) in the southwest, and Western Lake Erie Basin (WLEB) in the northwest) and across ten-year intervals such that patterns of community composition were assessed across 11 decades from the 1900s through the 2010s. First, descriptive patterns of total and aphidophagous lady beetle taxonomic beta-diversity were evaluated within geographic regions among decades by calculating total Sorenson dissimilarity (β_SOR_) using the *beta.temp* function in the R package ‘betapart’ (Baselga & Orme, 2012). Total Sørensen dissimilarity was decomposed into a turnover component (β_SIM_; reflects species replacement) and a nestedness component (β_SNF_; reflects species loss or gain) (Baselga, 2010). Next, pairwise dissimilarity matrices for Sørensen dissimilarity (β_sor_) and the respective components of species turnover (β_sim_) and nestedness (β_sne_) were calculated using the *beta.pair* function in the R package ‘betapart’ (Baselga & Orme, 2012). Then, permutational multivariate analysis of variance (PERMANOVA) and analysis of multivariate homogeneity of group dispersions (BETADISPER) were used to compare β_sor_, β_sim_, and β_sne_ pairwise dissimilarity matrices for total lady beetles and aphidophagous lady beetles among decades. PERMANOVA tests whether the centroid of communities differs among groups in multivariate space, while BETADISPER tests whether groups differ in the amount of dispersion from its spatial median among communities within a group. Differences in total and aphidophagous lady beetle community composition among decades were visualized using non-metric multidimensional scaling (NMDS). PERMANOVA, BETADISPER, and NMDS analyses were conducted using the ‘vegan’ package (Oksanen *et al*., 2011).

To determine the relative influence of temporal, landscape, spatial, and community drivers on the captures of five key species (*A. bipunctata, C. novemnotata, C. maculata, H. convergens*, the four most abundant native aphidophagous species, and *C. stigma*, the most abundant coccidophagous species), we constructed negative binomial generalized additive models (GAMs) to describe spatial and temporal patterns for each species using the ‘mgcv’ package (Wood, 2017), and then used an adaptive model selection procedure for each species in combination with available contextual data to determine the relative importance of the drivers. Response data took the form of total captures of each species in a given county, in a given decade. Because the absolute number of records was very sparse earlier in our study period, we restricted these analyses to specimens captured in or after 1930 for these analyses. For *A. bipunctata* and *C. novemnotata*, which were extremely rare or absent from collections in later decades, data were culled at 1990 to restrict these analyses of drivers to times when these species were present. We computed several community variables for each decade-county combination; in addition to absolute captures of each coccinellid species, including the abundance of two dominant invaders, *C. septempunctata* and *H. axyridis*, the total lady beetles captured, total alien species captured, and the proportion of the community captures belonging to alien species. All GAMs constructed included an offset variable of the structure log(1 + total lady beetles captured) to account for variability in sampling effort (with the exception of the model describing sampling effort over time).

First, to describe the relative abundance of each key species over time, corrected for varied sampling intensity, a simple GAM was constructed for each species with decade of capture as the independent variable, constrained to 3 knots and a smoothing parameter of 0.5. For the spatial analyses, data were aggregated into three-decade groups, and the captures of each species were modeled using a negative binomial GAM with a gaussian process smooth and a combination of latitude and longitude, both in aggregate and then by decade group. Latitude and longitude were included in the model to control for differences in the spatial distribution of lady beetle collections within the state and to account for spatial autocorrelation. To examine the interaction of temporal, spatial, landscape, and community factors on captures of each species, we used a modified backwards-stepping model selection applied to negative binomial GAMs. First, a global model was constructed that included decade, longitude, latitude, percentage land cover in agriculture, forest, and developed uses, the total captures of alien species, and the sampling offset. Because of a high-degree of autocorrelation between the aggregate alien species metrics, each of these variables was substituted into the model separately, and the variables with the best performance (determined by AIC) were selected for further analysis. Absolute numbers of *C. septempunctata* and *H. axyridis* were considered together in the same model to test if target species were exhibiting differential responses to the two alien species. After the substitution-based model selection was completed, the remaining model selection was completed using backwards selection by systematically dropping each variable with the lowest explanatory power, using AIC as the decision criterion. The selected model was then subjected to concurvity analysis and if the ‘worst case’ concurvity estimate exceeded 0.8 for any parameter, the parameter was eliminated from the model and backwards model selection was resumed (Ross, 2019). Complete data manipulation, community analyses, model specifications, model selection, and construction of prediction intervals, as well as the development history of our analyses are available on Github: https://github.com/BahlaiLab/Ohio_ladybeetles

## Results

We compiled 4,194 lady beetle museum records representing 28 species collected in Ohio, USA from 1900-2018 (Table 1; see Appendix 2: Table S2.3, Figs. S2.1-S2.4). Total collections of native species varied from year to year, with high numbers collected in the 1930s and 1980s (Fig. 3A). The most common native species represented in these collections were *Coleomegilla maculata* (Degeer) (16.4% of total records), *Hippodamia convergens* Guerin (9.5%), *Hippodamia parenthesis* (Say) (8.4%), *Psyllobora vigintimaculata* (Say) (7.6%), *Cycloneda munda* (Say) (7.2%), *Brachiacantha ursina* (Fabricius) (6.1%), and *Adalia bipunctata* (Linnaeus) (5.2%). Records documented the presence of the aphidophagous alien species *Coccinella septempunctata* (Linnaeus) (2.4% of total records; first detected in 1978), *Coccinella undecimpunctata* Linnaeus (0.02%; single record in 1953), *Harmonia axyridis* (Pallas) (11.2%; first detected in 1993), *Hippodamia variegata* (Goeze) (0.2%; first detected in 2000), and *Propylea quatuordecimpunctata* (Linnaeus) (0.6%; first detected in 2003). Collections of alien species began to increase in the 1980s, with numbers surpassing natives in the 2000s and 2010s (Fig. 3A).

**Figure 3.**
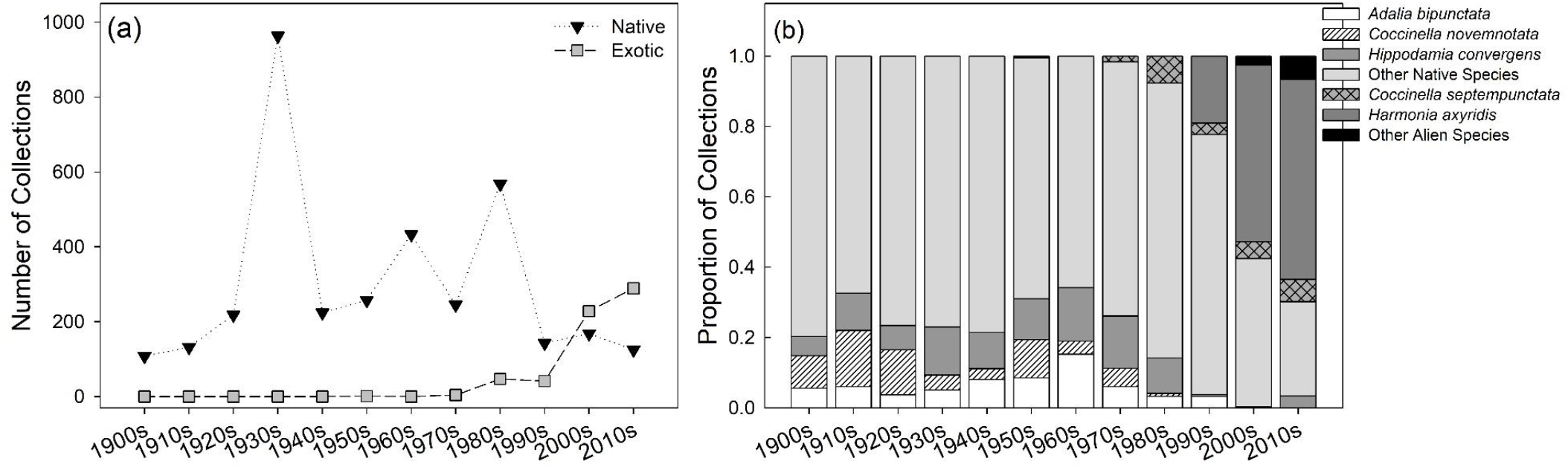
Number of collections (A) and proportion of collections (B) of native and alien lady beetle records from Ohio by decade from 1900 to 2018.

**Table 1.** Ohio lady beetle species records from 1900-2018 compiled from 25 institutions across the United States. Data collection focused on native and alien species within the tribe Coccinellini and four additional species (*B. ursina, C. stigma, H. undulata*, and *P. vigintimaculata*). Lady beetle species were characterized based on their status (native or alien to Ohio, USA) and their primary diet (aphids, scales, or fungi).

| Lady beetle species | Records | First Record | Last Record | Status | Primary Diet |
| --- | --- | --- | --- | --- | --- |
| *Brachiacantha ursina* (Fabricius) | 255 | 1902 | 2015 | Native | Aphidoidea, Coccoidea |
| *Hyperaspis undulata* (Say) | 61 | 1905 | 2014 | Native | Coccoidea |
| *Chilocorus stigma* (Say) | 111 | 1901 | 2016 | Native | Coccoidea, Aphidoidea |
| *Adalia bipunctata* (Linnaeus) | 218 | 1901 | 1996 | Native | Aphidoidea |
| *Anatis labiculata* (Say) | 176 | 1907 | 2012 | Native | Coccoidea, Aphidoidea |
| *Anatis mali* (Say) | 23 | 1932 | 2013 | Native | Coccoidea, Aphidoidea |
| *Anisosticta bitriangularis* (Say) | 16 | 1923 | 2012 | Native | Aphidoidea, Pollen |
| *Coccinella novemnotata* Herbst | 169 | 1901 | 1985 | Native | Aphidoidea |
| *Coccinella septempunctata* (Linnaeus) | 102 | 1978 | 2017 | Alien | Aphidoidea |
| *Coccinella transversoguttata* Mulsant | 51 | 1921 | 1986 | Native | Aphidoidea |
| *Coccinella trifasciata* Linnaeus | 19 | 1911 | 1971 | Native | Aphidoidea |
| *Coccinella undecimpunctata* Linnaeus | 1 | 1953 | 1953 | Alien | Aphidoidea |
| *Coleomegilla maculata* (Degeer) | 690 | 1901 | 2018 | Native | Aphidoidea, Pollen |
| *Cycloneda munda* (Say) | 302 | 1901 | 2017 | Native | Aphidoidea |
| *Harmonia axyridis* (Pallas) | 470 | 1993 | 2018 | Alien | Aphidoidea |
| *Hippodamia convergens* Guerin | 400 | 1903 | 2015 | Native | Aphidoidea |
| *Hippodamia glacialis* (Fabricius) | 39 | 1905 | 2016 | Native | Aphidoidea |
| *Hippodamia parenthesis* (Say) | 353 | 1900 | 2017 | Native | Aphidoidea |
| *Hippodamia quindecimmaculata* Mulsant | 3 | 1905 | 1935 | Native | Aphidoidea |
| *Hippodamia tredecimpunctata* (Linnaeus) | 155 | 1903 | 2014 | Native | Aphidoidea |
| *Hippodamia variegata* (Goeze) | 10 | 2000 | 2016 | Alien | Aphidoidea |
| *Mulsantina luteodorsa* J. Chapin | 1 | 2008 | 2008 | Native | Aphidoidea |
| *Mulsantina picta* (Randall) | 51 | 1924 | 2015 | Native | Aphidoidea |
| *Myzia pullata* (Say) | 30 | 1934 | 2012 | Native | Aphidoidea |
| *Neoharmonia venusta* (Melsheimer) | 119 | 1902 | 2012 | Native | Coccoidea |
| *Olla v-nigrum* (Mulsant) | 21 | 1932 | 2004 | Native | Aphidoidea, Psylloidea |
| *Propylea quatuordecimpunctata* (Linnaeus) | 27 | 2003 | 2016 | Alien | Aphidoidea |
| *Psyllobora vigintimaculata* (Say) | 321 | 1902 | 2016 | Native | Fungi (Erisyphaceae) |

### Lady beetle species composition

The proportion of native lady beetles comprising collections, including *A. bipunctata, C. novemnotata*, and *H. convergens*, has decreased in Ohio since the 1970s as alien lady beetles such as *C. septempunctata* and *H. axyridis* have increased (Fig. 3B). Descriptive patterns of lady beetle taxonomic beta-diversity (β_SOR_) within geographic regions across decades were the result of species turnover (β_SIM_) and nestedness (β_SNE_), but the strongest contributor to changes in lady beetle community composition shifted over time (Fig. 4A, B). Nestedness was the primary contributor to patterns of lady beetle beta-diversity until the 1980s. From 1980-2018, species turnover increasingly became the stronger contributor, especially for aphidophagous species, and was the dominant driver of lady beetle beta-diversity by the 2010s (Fig. 4A, B). PERMANOVA and NMDS analyses of pairwise beta-diversity matrices identified decades when significant shifts occurred in lady beetle community composition, primarily driven by aphidophagous species (see Appendix 2: Table S2.4, Fig. S2.5). Lady beetle beta-diversity (β_sor_) differed between the 1920s and 1930s (*F* = 3.88; *P* = 0.014) and between the 1930s and 1940s (*F* = 3.12; *P* = 0.003), with species composition in the 1930s being highly similar across geographic regions in Ohio and nested within the more variable collections from the 1920s and 1940s (see Appendix 2: Fig. S2.5A, B). Lady beetle beta-diversity (β_sor_) also differed between the 1980s and 1990s (*F* = 2.94; *P* = 0.021) and between the 1990s and 2000s (*F* = 3.36; *P* = 0.009), with communities similarly variable among decades, but collections shifting in species composition (see Appendix 2: Fig. S2.5C, D).

**Figure 4.**
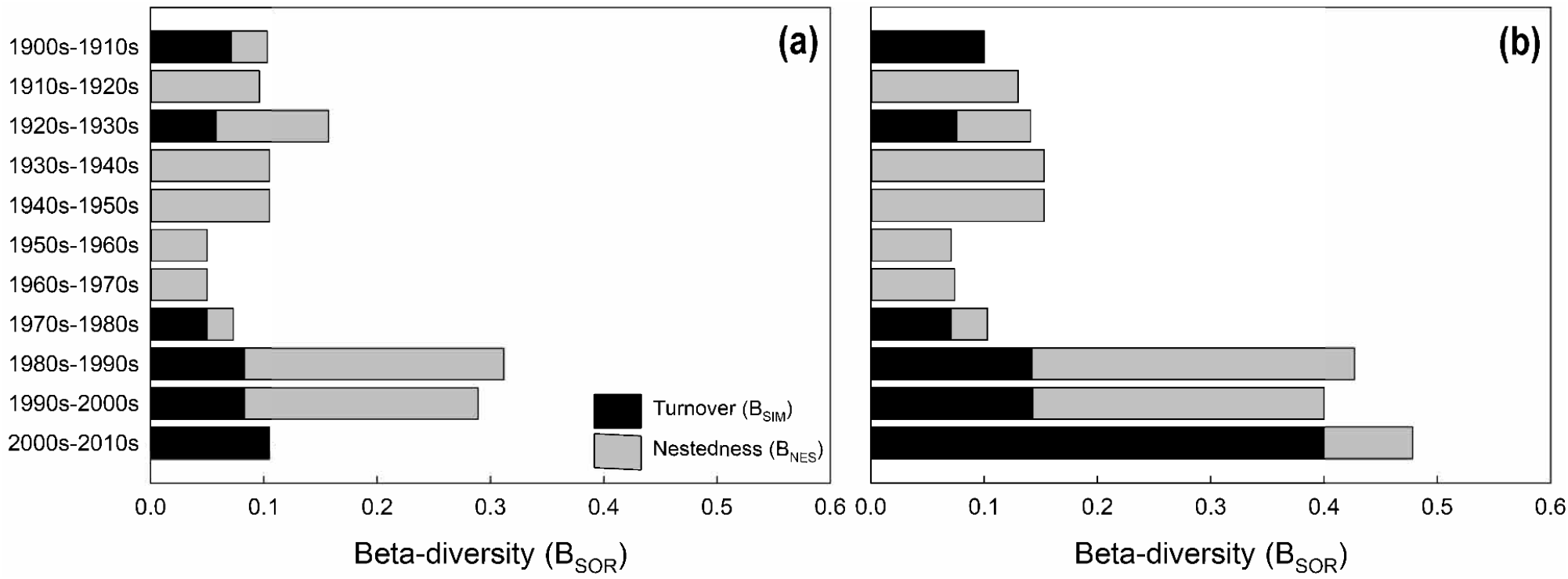
Descriptive patterns of taxonomic beta-diversity across decades for all lady beetle species (A) and aphidophagous lady beetle species (B) collected in Ohio. Total Sorensen dissimilarity (β_SOR_) among decades was partitioned into two additive components: turnover (β_SIM_; reflects species replacement) and nestedness (β_SNE_; reflects species loss or gain). Therefore, β_SOR_ = β_SIM_ + β_SNE_.

### Drivers of lady beetle populations

Relative captures of all lady beetle species varied with time, indicating that all species had some form of temporal dependence in the number of individuals captured when accounting only for sampling effort and using a normal error structure. However, these simple negative binomial models explained limited variation in the data. Using these simple models, native species were captured less per sampling effort later in the study period, but the steepness and timing of declines in captures varied by species. The two alien species exhibited more variable patterns. Captures of *C. septempuncata* initially increased in residual captures but then decreased during the four decades it has been present in Ohio. Conversely, captures of *H. axyridis* increased over the three decades since its establishment. For *A. bipunctata* and *C. novemnotata*, statistically significant temporal patterns were not observed within the modeled time period, as both species had relatively stable capture frequency prior to the 1980s but became extremely rare and then absent in later years of data collection. To provide meaningful model fits for the time when these species were present, years with zero-biased data were not included in their respective models. Sampling effort varied dramatically by year and location. Patterns of captures over time were not spatially static, as several native species also exhibited spatiotemporal dependencies over the study (i.e. changing spatial distributions with time; see Appendix 3: Table S3.1, Fig. S3.1-S3.5).

Species-specific patterns emerged in the negative binomial models that accounted for landscape, invasion, and spatiotemporal drivers, and model fit was much improved in all cases (Fig. 5; see Appendix 3: Table S3.1). *Adalia bipunctata* exhibited a steep negative response to the increasing proportion of alien species in the community but also was positively associated with agriculture and developed land covers. *Coccinella novemnotata* had a strong negative population trend over time as well as spatial dependencies in its captures but only appeared to respond negatively to agricultural cover as a landscape driver. Since this native species had already begun to decline prior to invasions by *C. septempunctata* and *H. axyridis*, there was limited co-occurrence of *C. novemnotata* and any of the alien species (only 5 captures of *C. novemnotata* were recorded after 1980). *Coleomegilla maculata* populations were relatively net stable over time, with an increase peaking in the 1970s and 1980s, but a net decrease since. Our models suggest a relatively neutral effect of small numbers of alien lady beetles on *C. maculata*, but a negative impact as alien species become dominant (i.e. >50% of the lady beetles collected). Additionally, agricultural land cover had a slight positive association with captures of *C. maculata*. Captures of *H. convergens* had a negative trend over time and exhibited a differential response to the two dominant alien species: a positive association with *C. septempunctata* and a negative association with *H. axyridis*, as well as a positive association with higher values of developed land cover. *Chilocorus stigma* exhibited spatial dependencies and a positive association with forested habitats, and a negative association with the proportion of alien lady beetles in the community.

**Figure 5.**
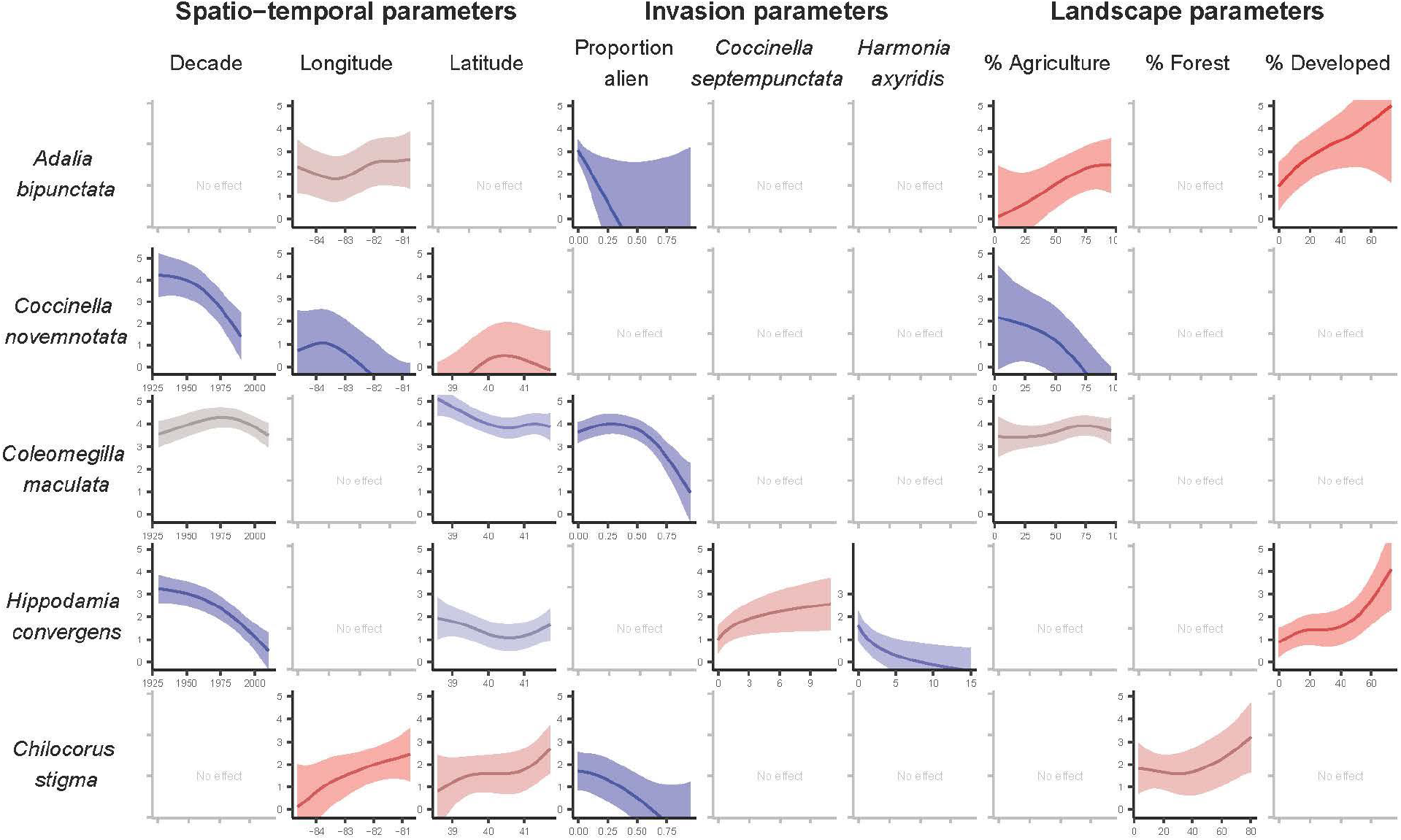
Matrix of partial GAM predictions of responses for five key native species to spatiotemporal, invasion, and landscape parameters. Negative binomial generalized additive models used Ohio museum collections, 1930-2018 data on lady beetle captures and were adjusted for sampling effort. Predictions were constructed by generating data that held all other parameters constant and varying the parameter of interest, and then substituting these data into the best-fit model determined by model selection (Appendix 2, Table 1). Solid lines, shading represent 95% confidence intervals.

## Discussion

Specimen records can be harnessed to understand long-term biodiversity trends across various spatial scales in response to anthropogenic threats (Shaffer *et al*., 1998; Suarez & Tsutsui, 2004; Lister, 2011; Meineke *et al*., 2019). Using historic occurrence records of lady beetles collected in Ohio, USA, we report evidence of declines in captures for several native species, but the timing and severity of declines as well as the relative importance of alien species and land cover change as long-term drivers were species-specific. Observed changes in species composition that began in the 1980s indicated processes of species loss/gain and turnover that shifted communities towards dominance of a few alien lady beetle species.

### Shifts in lady beetle species composition

As collections of some native species declined over time in Ohio, captures of alien lady beetle species increased following their establishment and spread. Beginning in the 2000s, alien species comprised over 60% of the total specimens collected, suggesting a shift in lady beetle community structure and alien species dominance within the state. This shift was further supported by changes in patterns of total beta-diversity within geographic regions of the state that began in the 1980s. From the 1980s to the 2010s, beta-diversity of lady beetle species increased across decades, and the contribution of species turnover to these patterns increased compared to previous decades, becoming the dominant driver of beta-diversity by the 2010s. This finding indicates that lady beetle communities within a geographic region became increasingly dissimilar over time due to processes of species loss or gain (i.e. species nestedness) and species replacement (i.e. species turnover) during the period that coincides with alien lady beetle establishment and spread. Changes in lady beetle species composition from the 1900s to the 1970s were primarily driven by species nestedness. These patterns were the result of the loss and gain of uncommon native species in collections throughout the state, as well as the loss of common native species such as *A. bipunctata* and *C. novemnotata* which began to decline prior to alien species arrival. Species composition shifted in the 1980s as dissimilarity of communities increased substantially and the contribution of species replacement to these patterns increased. High species replacement indicates that a similar number of lady beetle species were collected within geographic regions among these decades, but a low number were shared as species were replaced over time.

A combination of species nestedness and species turnover that resulted in high dissimilarity in lady beetle species composition during the 1980s, 1990s, and 2000s is indicative of spatiotemporal loss and replacement of native species. By the 2010s, lady beetle communities had become more similar to each other in terms of species composition and were dominated by alien lady beetle species. Similar shifts in lady beetle communities have been observed in response to the establishment of alien species (Elliott *et al*., 1996; Alyokhin & Sewell, 2004; Bahlai *et al*., 2013). For example, a shift in species dominance with the establishment of *C. septempunctata* in the 1980s was observed in potato fields in Maine, USA (Alyokhin & Sewell, 2004). Within four years of its establishment in southwestern Michigan, USA, *H. axyridis* was the dominant lady beetle species collected and was found in diverse habitats, including agricultural and old fields, and poplar plantations (Colunga-Garcia & Gage, 1998). Although the composition of lady beetle communities has become dominated by exotic species, Bahlai et al. (2013) documented that their potential to suppress pest populations in agricultural and natural habitats had remained relatively consistent over time. As significant shifts in lady beetle communities have occurred across the Midwestern US, reliance on alien species to maintain pest suppression may be required for management, but further declines of native species may affect long-term resilience of this ecosystem service (Bahlai *et al*., 2021).

### Drivers of native lady beetle decline

Using museum specimens collected over twelve decades, we documented declines in captures of the native species *A. bipunctata, C. novemnotata, H. convergens, C. maculata*, and *C. stigma*. The native species *A. bipunctata, C. novemnotata*, and *H. convergens* were once widely distributed across much of the United States (Gordon, 1985), with early surveys in Ohio recording these species across the state (Dury, 1879; Bubna, 1902). Now, these species are considered rare or potentially extirpated in much of eastern North America (Wheeler Jr & Hoebeke, 1995; Elliott *et al*., 1996; Gardiner *et al*., 2012). For example, no captures of *C. novemnotata* or *A. bipunctata* were found after 1985 and 1996, respectively, suggesting their populations were below the detection threshold or extirpated within the state. While declines in captures of these native species were observed, the relative importance of alien lady beetles and land cover change as drivers was species-specific.

### Adalia bipunctata

We found evidence that declines in captures of the aphidophagous species *A. bipunctata* began in the 1960s, prior to the arrival and dominance of alien lady beetle species. The last collection of *A. bipunctata* in Ohio occurred in 1996, and since the arrival of *C. septempunctata* in 1978, there were only 13 individuals of this native species collected from the state. During this period of overlap with alien species, collections of *A. bipunctata* decreased as the proportion of captures of alien species increased within the community. Reports from southwestern Michigan also observed declines in *A. bipunctata* over 24 years with increasing dominance of alien species (Bahlai *et al*., 2015). *Harmonia axyridis* was first collected in Ohio in 1993 and shares an affinity for arboreal habitats with *A. bipunctata* (Colunga-Garcia & Gage, 1998; Honek *et al*., 2019). Although our findings indicate that alien species likely contributed to declines in captures of *A. bipunctata* to some degree, declines in captures prior to the arrival of alien species suggests other contributing factors such as landscape change. We found that collections of *A. bipunctata* increased with percentage of agricultural and developed land cover. Land cover change analyses showed agricultural land cover decreased, while developed land cover increased over time. We hypothesize that declines in captures of this native species could be related to reductions in aphid prey availability due to a combination of landscape stressors such as reductions in agricultural land cover paired with improved pest management practices as well as improved air and/or habitat quality in urban environments that reduce pest outbreaks (Sloggett, 2017).

### Coccinella novemnotata

We found no evidence that declines in captures of the aphidophagous species *C. novemnotata* in Ohio were related to alien lady beetles, as this species was nearly absent from collections by the time *C. septempunctata* was recorded in 1978. Captures of *C. novemnotata* began to decline in the 1950s prior to the arrival of alien species, with the last Ohio collection occurring in 1985. Since the arrival of *C. septempuncata* in Ohio, only 12 individuals of this native species were recorded. This finding contrasts with many studies that have hypothesized declines of *C. novemnotata* were related to competitive displacement by *C. septempunctata* (Staines Jr *et al*., 1990; Wheeler Jr & Hoebeke, 1995; Snyder & Evans, 2006). For instance, declines of *C. novemnotata* have been widely reported during the 1970s and 1980s, which coincides with the establishment and spread of *C. septempunctata* (Wheeler Jr & Hobeke, 1995 and references therein). We found that captures of *C. novemnotata* decreased with increasing agricultural land cover, with declines beginning around the 1950s. Since the 1930s, the amount of agricultural land cover has decreased within the state, but the spatial extent of these changes was highly variable and not consistent across all Ohio counties. This loss of agriculture was primarily in the eastern region of the state, while western Ohio remained dominated by agricultural land cover throughout the study period. Along with these changes in the extent of agricultural land cover, landscapes have experienced a shift from more diversified cropping systems to highly managed crop monocultures since the mid-1900s (Crossley *et al*., 2021). *Coccinella novemnotata* has a broad ecological niche (Losey *et al*., 2007), and populations have been found in a variety of cultivated crops (Wheeler Jr & Hoebeke, 1995), including alfalfa (Goodarzy & Davis, 1958; Pimentel & Wheeler, 1973), corn (Smith, 1971), cotton (Bell & Whitcomb, 1964), soybeans (Dumas *et al*., 1964), and fruit trees (Oatman *et al*., 1964; Putman, 1964). Simplification of agricultural landscapes due to loss of natural habitat and reductions in crop diversity may impact temporal prey availability and diversity as well as refuge and overwintering habitats that are required for predatory insect species such as *C. novemnotata* (Rusch *et al*., 2016).

### Coleomegilla maculata

Captures of the aphidophagous species *C. maculata* increased slightly and then declined with increasing dominance of alien species within the community. *Coleomegilla maculata* is known to be a strong aphidophagous competitor (Long & Finke, 2014), and this species remains one of the most frequently collected native lady beetles in croplands and grasslands (Smith and Gardiner 2013). For example, *C. maculata* was commonly collected in residential gardens across the state of Ohio by community scientists (Gardiner *et al*., 2012; Gardiner *et al*., 2021). However, declines in *C. maculata* populations were reported during a long-term study in southwestern Michigan (Bahlai *et al*., 2015). *Coleomegilla maculata* feeds on pollen in addition to aphid prey (Dixon & Dixon, 2000; Hodek *et al*., 2012; Majerus, 2016), and this pollen feeding was hypothesized to reduce competition with alien species (Bahlai *et al*., 2015). However, there is evidence that *H. axyridis* and *C. septempunctata* also feed on pollen as a supplemental food resource when arthropod prey is scarce (Ricci *et al*., 2005; Berkvens *et al*., 2008; Berkvens *et al*., 2010). Therefore, the dietary niche overlap and thus the impacts of alien species on *C. maculata* may have been underestimated.

### Hippodamia convergens

Our findings isolated *H. axyridis* as the major driver of declines in captures of the aphidophagous species *H. convergens*. *Hippodamia convergens* was last collected in Ohio in 2015, although only 18 individuals are recorded from 1990-2015. Collections of *H. convergens* declined with captures of *H. axyridis* but increased with captures of *C. septempunctata*. These divergent patterns highlight that alien lady beetle species are not ecologically equivalent, and instead, can have different impacts on native species. *Hippodamia convergens* also was positively associated with percentage of developed land. The effects of urbanization on lady beetles are highly context dependent, as previous studies have reported positive (Egerer *et al*., 2017; Honek *et al*., 2018; Liere *et al*., 2019) and negative (Rocha *et al*., 2018; Grez *et al*., 2019; Parker *et al*., 2020) associations. However, as *H. convergens* has widely declined within the study region over the past several decades while urban habitat has increased, this observed positive association does not appear to be a major driver of their population status and could reflect a sampling bias wherein this rare species was more likely to be collected in densely populated areas.

### Chilocorus stigma

Collections of the coccidophagous species *C. stigma* declined as the proportion of alien species increased in the community, suggesting that aphidophagous alien lady beetles may affect non-aphidophagous native species to some extent. Likewise, decline of *C. stigma* immediately following the establishment of *H. axyridis* has been reported in southwestern Michigan, USA, further suggesting potential competitive interactions among these species (Colunga-Garcia & Gage, 1998). Although *H. axyridis* is primarily aphidophagous (Koch, 2003), this species also feeds on scale insects in arboreal environments (Mcclure, 1986), which suggests some degree of dietary and habitat overlap with *C. stigma. Chilocorus stigma* was the only native lady beetle associated with forest land cover, with captures increasing as the percentage of forest increased at the county level. This response is likely linked with the ecology of this species. Scale insects are common pests of trees in managed and natural forests. Further, *C. stigma* is reported to oviposit eggs in bark cracks and crevices as well as overwinter as adults in the leaf litter layer (Mayer & Allen, 1983). Similar patterns have been found in other native lady beetle surveys. For instance, the amount of forest habitat at a 2 km landscape scale was positively associated with native lady beetle abundance and species diversity within residential gardens (Gardiner *et al*., 2021). The conservation value of forests for native lady beetles may be underestimated, but additional research is required to understand when and how native species are utilizing these habitats.

### Conclusions

Using specimen records collected over 118 years in Ohio, USA, we documented shifts in lady beetle species composition beginning in the 1980s as communities became increasingly dominated by alien species. Because of uneven sampling inherent to museum collections, total records of native lady beetles varied from year to year, which made it difficult to detect any changes in absolute abundance at the community-level. Therefore, our methodology and results cannot provide evidence that all native lady beetles are in decline. Such stochasticity may be inherent to lady beetle biology as native populations are known to experience boom-bust cycles (Bahlai *et al*., 2015) and other studies have reported similar year to year variability in their numbers (Elliott & Kieckhefer, 1990; Kieckhefer & Elliott, 1990; Elliott *et al*., 1996; Harmon *et al*., 2007). Despite patterns of annual variation, we detected evidence of declines in captures of several native lady beetle species, including several aphidophagous species and a coccidophagous species, via decreasing representation in the sampled community of lady beetles.

The use of long-term specimen records facilitated investigation of the relative importance of the establishment of alien species and landscape change as drivers of native lady beetle decline. Drivers of declines in captures of native lady beetles were highly species-specific, emphasizing that mechanisms driving population losses cannot be generalized even among closely related species. Additionally, this finding highlights the importance of species-level data when investigating temporal trends in insect populations. Although the establishment of exotic species and landscape change have been identified as major drivers of spatiotemporal patterns in insect populations (Fox *et al*., 2014; Sánchez-Bayo & Wyckhuys, 2019; Seibold *et al*., 2019), the causes of declines are likely more complex and multifaceted (Homburg *et al*., 2019; Wagner, 2020; Wagner *et al*., 2021). Our study underscores this complexity by documenting how closely related native lady beetle species displayed opposing, species-specific responses to alien species and land cover change. For several native species investigated, the dominance of alien lady beetles was identified as a major contributor to declines in captures, but other native species began to decline prior to alien species establishment. The importance of landscape change as a driver structuring the distributions of lady beetle populations suggests biodiversity conservation management is required at the landscape scale. Landscape scale management will need to balance the opposing needs of native species to be effective. Importantly, we are unable to disentangle the effects of historical changes in land cover with more recent intensification and simplification of agricultural landscapes on these observed trends in captures. Native lady beetle species are key predators of aphids, scales, psyllids, mites, fungi, and other pests (Evans, 2009; Hodek & Honěk, 2009; Weber & Lundgren, 2009), contributing broadly to biological control in agricultural systems (Caltagirone & Doutt, 1989; Obrycki & Kring, 1998; Rondoni *et al*., 2021). Increased dominance of alien lady beetles indicates that these species may be required to maintain successful pest management in the future. Understanding how these major anthropogenic drivers influence long-term trends in native lady beetle populations will inform the conservation of this ecologically and economically important family of insects.

## Acknowledgements

We acknowledge the institutions and their curators for their time and assistance with compiling lady beetle specimen records: California Academy of Sciences (Susan Gin and Christopher C. Grinter); Natural History Museum of Los Angeles County (Brian Brown and Weiping Xie); Peabody Museum, Yale University (Lawrence Gall); Biodiversity Research Collections, University of Connecticut (Jane O’Donnell); Smithsonian National Museum of Natural History (Natalie Vandenburgh); Collection of Arthropods, University of Georgia (Joseph V. McHugh and E. Richard Hoebeke); Gantz Family Collections Center, The Field Museum (Rebekah Shuman Baquiran); Illinois Natural History Survey, University of Illinois (Thomas McElrath); Louisiana State Arthropod Museum, Louisiana State University (Victoria Moseley Bayless); A.J. Cook Arthropod Research Collection, Michigan State University (Gary L. Parsons); Museum of Zoology, University of Michigan (Erika Tucker); University of Minnesota (Robin Thomson); Wilbur R. Enns Entomology Museum, University of Missouri (Kristin B. Simpson); Mississippi Entomological Museum, Mississippi State University (Terence L. Schiefer); C.A. Triplehorn Insect Collection, The Ohio State University (Luciana Musetti); Cleveland Museum of Natural History (Nicole Gunter); Carnegie Museum of Natural History (Robert Androw and Robert Davidson); The Frost Entomological Museum, Pennsylvania State University; Clemson University Arthropod Collection, Clemson University (Michael Ferro); Severin-McDaniel Insect Research Collection, South Dakota State University (Louis Hesler); Brigham Young University (Shawn M. Clark); Crane Hollow Nature Preserve (Gary and Holly Coovert); Agricultural Technical Institute Teaching Collection, The Ohio State University (Jon Van Gray). Ohio lady beetle records from Boonshoft Museum of Discovery were provided by Kaitlin U. Campbell and Thomas O. Crist, and records from The Lost Ladybug Project (http://www.lostladybug.org/) were made available by Rebecca Smyth. This work was partially supported by a grant to CAB from the National Science Foundation DBI #2045721.

## Author Contributions

M.M. Gardiner, K.I. Perry, and C.A. Bahlai conceived, designed, and implemented the study; K.I. Perry, C.B. Riley, K. J. Turo, L. Taylor, J. Radl, Y.A. Delegado de la flor, and F. S. Sivakoff compiled the specimen data; K.I. Perry, C.A. Bahlai, and T.J. Assal analyzed and interpreted the data; K.I. Perry wrote the first draft of the manuscript; all authors reviewed and edited the manuscript.

## Data availability statement

The data and analyses code that support the findings of this study are openly available on GitHub: https://github.com/BahlaiLab/Ohio_ladybeetles

## References

Alyokhin, A. & Sewell, G. (2004) Changes in a lady beetle community following the establishment of three alien species. Biological Invasions, 6, 463–471.

Angalet, G.W., Tropp, J.M. & Eggert, A.N. (1979) *Coccinella septempunctata* in the United States: Recolonizations and notes on its ecology. Environmental Entomology, 8, 896–901.

Bahlai, C.A. & Zipkin, E.F. (2020) The Dynamic Shift Detector: An algorithm to identify changes in parameter values governing populations. PLOS Computational Biology, 16, e1007542.

Bahlai, C.A., Colunga-Garcia, M., Gage, S.H. & Landis, D.A. (2013) Long-yerm functional dynamics of an aphidophagous coccinellid community remain unchanged despite repeated invasions. PLoS ONE, 8, e83407.

Bahlai, C.A., Colunga-Garcia, M., Gage, S.H. & Landis, D.A. (2015) The role of exotic lady beetles in the decline of native lady beetle populations: Evidence from long-term monitoring. Biological Invasions, 17, 1005–1024.

Bahlai, C.A., Hart, C., Kavanaugh, M.T., White, J.D., Ruess, R.W., Brinkman, T.J., Ducklow, H.W., Foster, D.R., Fraser, W.R., Genet, H., Groffman, P.M., Hamilton, S.K., Johnstone, J.F., Kielland, K., Landis, D.A., Mack, M.C., Sarnelle, O. & Thompson, J.R. (2021) Cascading effects: insights from the U.S. Long Term Ecological Research Network. Ecosphere, 12, e03430.

Baselga, A. (2010) Partitioning the turnover and nestedness components of beta diversity. Global Ecology and Biogeography, 19, 134–143.

Baselga, A. & Orme, C.D.L. (2012) betapart: an R package for the study of beta diversity. Methods in Ecology and Evolution, 3, 808–812.

Bell, K.O. & Whitcomb, W.H. (1964) Field studies on egg predators of the bollworm, *Heliothis zea* (Boddie). The Florida Entomologist, 47, 171–180.

Berkvens, N., Bonte, J., Berkvens, D., Deforce, K., Tirry, L. & De Clercq, P. (2008) Pollen as an alternative food for *Harmonia axyridis*. BioControl, 53, 201–210.

Berkvens, N., Landuyt, C., Deforce, K., Berkvens, D., Tirry, L. & De Clercq, P. (2010) Alternative foods for the multicoloured Asian lady beetle *Harmonia axyridis* (Coleoptera: Coccinellidae). European Journal of Entomology, 107, 189–195.

Boakes, E.H., McGowan, P.J.K., Fuller, R.A., Chang-qing, D., Clark, N.E., O’Connor, K. & Mace, G.M. (2010) Distorted views of biodiversity: Spatial and temporal bias in species occurrence data. PLOS Biology, 8, e1000385.

Brooks, D.R., Bater, J.E., Clark, S.J., Monteith, D.T., Andrews, C., Corbett, S.J., Beaumont, D.A. & Chapman, J.W. (2012) Large carabid beetle declines in a United Kingdom monitoring network increases evidence for a widespread loss in insect biodiversity. Journal of Applied Ecology, 49, 1009–1019.

Brown, P.M.J. & Roy, H.E. (2018) Native ladybird decline caused by the invasive harlequin ladybird *Harmonia axyridis*: Evidence from a long-term field study. Insect Conservation and Diversity, 11, 230–239.

Brown, P.M.J., Ingels, B., Wheatley, A., Rhule, E.L., de Clercq, P., van Leeuwen, T. & Thomas, A. (2015) Intraguild predation by *Harmonia axyridis* (Coleoptera: Coccinellidae) on native insects in Europe: Molecular detection from field samples. Entomological Science, 18, 130–133.

Bubna, M. (1902) Coleoptera of Cuyahoga County, Ohio. Ohio Naturalist, 2, 193–197.

Caltagirone, L.E. & Doutt, R.L. (1989) The history of the Vedalia beetle importation to California and its impact on the development of biological control. Annual Review of Entomology, 34, 1–16.

Cardoso, P., Barton, P.S., Birkhofer, K., Chichorro, F., Deacon, C., Fartmann, T., Fukushima, C.S., Gaigher, R., Habel, J.C., Hallmann, C.A., Hill, M.J., Hochkirch, A., Kwak, M.L., Mammola, S., Ari Noriega, J., Orfinger, A.B., Pedraza, F., Pryke, J.S., Roque, F.O., Settele, J., Simaika, J.P., Stork, N.E., Suhling, F., Vorster, C. & Samways, M.J. (2020) Scientists’ warning to humanity on insect extinctions. Biological Conservation, 242, 108426.

Ceballos, G., Ehrlich, P.R., Barnosky, A.D., García, A., Pringle, R.M. & Palmer, T.M. (2015) Accelerated modern human-induced species losses: Entering the sixth mass extinction. Science Advances, 1, e1400253.

Chen, I.-C., Shiu, H.-J., Benedick, S., Holloway, J.D., Chey, V.K., Barlow, H.S., Hill, J.K. & Thomas, D. (2009) Elevation increases in moth assemblages over 42 years on a tropical mountain. Proceedings of the National Academy of Sciences, 106, 1479–1483.

Colla, S.R., Gadallah, F., Richardson, L., Wagner, D. & Gall, L. (2012) Assessing declines of North American bumble bees *(Bombus* spp.) using museum specimens. Biodiversity and Conservation, 21, 3585–3595.

Colunga-Garcia, M. & Gage, S.H. (1998) Arrival, establishment, and habitat use of the multicolored Asian lady beetle (Coleoptera: Coccinellidae) in a Michigan landscape. Environmental Entomology, 27, 1574–1580.

Conrad, K.F., Warren, M.S., Fox, R., Parsons, M.S. & Woiwod, I.P. (2006) Rapid declines of common, widespread British moths provide evidence of an insect biodiversity crisis. Biological Conservation, 132, 279–291.

Crossley, M.S., Burke, K.D., Schoville, S.D. & Radeloff, V.C. (2021) Recent collapse of crop belts and declining diversity of US agriculture since 1840. Global Change Biology, 27, 151–164.

Crossley, M.S., Meier, A.R., Baldwin, E.M., Berry, L.L., Crenshaw, L.C., Hartman, G.L., Lagos-Kutz,, Nichols, D.H., Patel, K., Varriano, S., Snyder, W.E. & Moran, M.D. (2020) No net insect abundance and diversity declines across US Long Term Ecological Research sites. Nature Ecology & Evolution, 4, 1368–1376.

Dewitz, J. (2019) National Land Cover Database (NLCD) 2016 Products (ver. 2.0, July 2020): U.S. Geological Survey data release. https://doi.org/10.5066/P96HHBIE.

Didham, R.K., Tylianakis, J.M., Gemmell, N.J., Rand, T.A. & Ewers, R.M. (2007) Interactive effects of habitat modification and species invasion on native species decline. Trends in Ecology & Evolution, 22, 489–496.

Didham, R.K., Basset, Y., Collins, C.M., Leather, S.R., Littlewood, N.A., Menz, M.H.M., Müller, J., Packer, L., Saunders, M.E., Schönrogge, K., Stewart, A.J.A., Yanoviak, S.P. & Hassall, C. (2020) Interpreting insect declines: Seven challenges and a way forward. Insect Conservation and Diversity, 13, 103–114.

Diepenbrock, L.M. & Finke, D.L. (2013) Refuge for native lady beetles (Coccinellidae) in perennial grassland habitats. Insect Conservation and Diversity, 6, 671–679.

Diepenbrock, L.M., Fothergill, K., Tindall, K.V., Losey, J.E., Smyth, R.R. & Finke, D.L. (2016) The influence of exotic lady beetle (Coleoptera: Coccinellidae) establishment on the species composition of the native lady beetle community in Missouri. Environmental Entomology, 45, 855–864.

Dirzo, R. & Raven, P.H. (2003) Global state of biodiversity and loss. Annual Review of Environment and Resources, 28, 137–167.

Dixon, A.F.G. & Dixon, A.E. (2000) Insect predator-prey dynamics: ladybird beetles and biological control. Cambridge University Press.

Dumas, B.A., Boyer, W.P. & Whitcomb, W.H. (1964) Effect of various factors on surveys of predaceous insects in soybeans. Journal of the Kansas Entomological Society, 37, 192–201.

Dury, C. (1879) List of the Coleoptera observed in the vicinity of Cincinnati. The Journal of the Cincinnati Society of Natural History, 2, 162–178.

Egerer, M., Li, K. & Ong, T.W. (2018) Context matters: Contrasting ladybird beetle responses to urban environments across two US regions. Sustainability, 10

Egerer, M.H., Bichier, P. & Philpott, S.M. (2017) Landscape and local habitat correlates of lady beetle abundance and species richness in urban agriculture. Annals of the Entomological Society of America, 110, 97–103.

Elliott, N.C. & Kieckhefer, R.W. (1990) Dynamics of aphidophagous coccinellid assemblages in small grain fields in eastern South Dakota. Environmental Entomology, 19, 1320–1329.

Elliott, N.C., Kieckhefer, R. & Kauffman, W. (1996) Effects of an invading coccinellid on native coccinellids in an agricultural landscape. Oecologia, 105, 537–544.

Entomological Collections Network (ENC) (2020) Collections and Archives. Retrieved from https://ecnweb.net/resources/collections/, 30 October 2020.

Evans, E.W. (2004) Habitat displacement of North American ladybirds by an introduced species. Ecology, 85, 637–647.

Evans, E.W. (2009) Lady beetles as predators of insects other than Hemiptera. Biological Control, 51, 255–267.

Fox, R., Oliver, T.H., Harrower, C., Parsons, M.S., Thomas, C.D. & Roy, D.B. (2014) Long-term changes to the frequency of occurrence of British moths are consistent with opposing and synergistic effects of climate and land-use changes. Journal of Applied Ecology, 51, 949–957.

Gagnon, A.-È., Heimpel, G.E. & Brodeur, J. (2011) The ubiquity of intraguild predation among predatory arthropods. PLoS ONE, 6, e28061.

Gardiner, M.M., Allee, L.L., Brown, P.M.J., Losey, J.E., Roy, H.E. & Smyth, R.R. (2012) Lessons from lady beetles: Accuracy of monitoring data from US and UK citizen-science programs. Frontiers in Ecology and the Environment, 10, 471–476.

Gardiner, M.M., Perry, K.I., Riley, C.B., Turo, K.J., Delgado de la flor, Y.A. & Sivakoff, F.S. (2021) Community science data suggests that urbanization and forest habitat loss threaten aphidophagous native lady beetles. Ecology and Evolution, 11, 2761–2774.

Gardiner, M.M., Landis, D.A., Gratton, C., Schmidt, N., O’Neal, M., Mueller, E., Chacon, J., Heimpel, G.E. & DiFonzo, C.D. (2009) Landscape composition influences patterns of native and exotic lady beetle abundance. Diversity and Distributions, 15, 554–564.

Goodarzy, K. & Davis, D.W. (1958) Natural enemies of the spotted alfalfa aphid in Utah. Journal of Economic Entomology, 51, 612–616.

Gordon, R.D. (1985) The Coccinellidae (Coleoptera) of America north of Mexico. Journal of the New York Entomological Society, 93, 912 pp.

Grez, A.A., Rand, T.A., Zaviezo, T. & Castillo-Serey, F. (2013) Land use intensification differentially benefits alien over native predators in agricultural landscape mosaics. Diversity and Distributions, 19, 749–759.

Grez, A.A., Zaviezo, T., Gardiner, M.M. & Alaniz, A.J. (2019) Urbanization filters coccinellids composition and functional trait distributions in greenspaces across greater Santiago, Chile. Urban Forestry & Urban Greening, 38, 337–345.

Grixti, J.C., Wong, L.T., Cameron, S.A. & Favret, C. (2009) Decline of bumble bees (*Bombus*) in the North American Midwest. Biological Conservation, 142, 75–84.

Habel, J.C., Segerer, A., Ulrich, W., Torchyk, O., Weisser, W.W. & Schmitt, T. (2016) Butterfly community shifts over two centuries. Conservation Biology, 30, 754–762.

Hallmann, C.A., Sorg, M., Jongejans, E., Siepel, H., Hofland, N., Schwan, H., Stenmans, W., Müller, A., Sumser, H., Hörren, T., Goulson, D. & de Kroon, H. (2017) More than 75 percent decline over 27 years in total flying insect biomass in protected areas. PLoS ONE, 12, e0185809.

Harmon, J.P., Stephens, E. & Losey, J. (2007) The decline of native coccinellids (Coleoptera: Coccinellidae) in the United States and Canada. Journal of Insect Conservation, 11, 85–94.

Hijmans, R.J. (2020) raster: Geographic data analysis and modeling, R package version 3.5-2, https://cran.r-project.org/web/packages/raster/index.html.

Hodek, I. & Honěk, A. (2009) Scale insects, mealybugs, whiteflies and psyllids (Hemiptera, Sternorrhyncha) as prey of ladybirds. Biological Control, 51, 232–243.

Hodek, I. & Michaud, J. (2013) Why is *Coccinella septempunctata* so successful? (A point-of-view). European Journal of Entomology, 105, 1–12.

Hodek, I., Honek, A. & Van Emden, H.F. (2012) Ecology and behaviour of the ladybird beetles (Coccinellidae). John Wiley & Sons.

Honek, A., Martinkova, Z., Roy, H.E., Dixon, A.F., Skuhrovec, J., Pekár, S. and Brabec, M. (2019) Differences in the phenology of *Harmonia axyridis* (Coleoptera: Coccinellidae) and native coccinellids in Central Europe. Environmental Entomology, 48, 80–87.

Homburg, K., Drees, C., Boutaud, E., Nolte, D., Schuett, W., Zumstein, P., von Ruschkowski, E. & Assmann, T. (2019) Where have all the beetles gone? Long-term study reveals carabid species decline in a nature reserve in Northern Germany. Insect Conservation and Diversity, 12, 268–277.

Honek, A., Martinkova, Z. & Strobach, J. (2018) Effect of aphid abundance and urbanization on the abundance of *Harmonia axyridis* (Coleoptera: Coccinellidae). European Journal of Entomology, 115, 703–707.

Honek, A., Martinkova, Z., Kindlmann, P., Ameixa, O.M.C.C. & Dixon, A.F.G. (2014) Long-term trends in the composition of aphidophagous coccinellid communities in Central Europe. Insect Conservation and Diversity, 7, 55–63.

Honek, A., Dixon, A.F.G., Soares, A.O., Skuhrovec, J. & Martinkova, Z. (2017) Spatial and temporal changes in the abundance and compostion of ladybird (Coleoptera: Coccinellidae) communities. Current Opinion in Insect Science, 20, 61–67.

Johnson, K.G., Brooks, S.J., Fenberg, P.B., Glover, A.G., James, K.E., Lister, A.M., Michel, E., Spencer, M., Todd, J.A., Valsami-Jones, E., Young, J.R. & Stewart, J.R. (2011) Climate change and biosphere response: Unlocking the collections vault. BioScience, 61, 147–153.

Katsanis, A., Babendreier, D., Nentwig, W. & Kenis, M. (2013) Intraguild predation between the invasive ladybird *Harmonia axyridis* and non-target European coccinellid species. BioControl, 58, 73–83.

Kharouba, H.M., Lewthwaite, J.M.M., Guralnick, R., Kerr, J.T. & Vellend, M. (2019) Using insect natural history collections to study global change impacts: challenges and opportunities. Philosophical Transactions of the Royal Society B: Biological Sciences, 374, 20170405.

Kieckhefer, R.W. & Elliott, N.C. (1990) A 13-year survey of the aphidophagous Coccinellidae in maize fields in eastern South Dakota. The Canadian Entomologist, 122, 579–581.

Koch, R.L. (2003) The multicolored Asian lady beetle, *Harmonia axyridis:* A review of its biology, uses in biological control, and non-target impacts. Journal of Insect Science, 3:32

LaMana, M.L. & Miller, J.C. (1996) Field observations on *Harmonia axyridis* Pallas (Coleoptera: Coccinellidae) in Oregon. Biological Control, 6, 232–237.

Li, H., Li, B., Lövei, G.L., Kring, T.J. & Obrycki, J.J. (2021) Interactions among native and non-native predatory Coccinellidae influence biological control and biodiversity. Annals of the Entomological Society of America, 114, 119–136.

Liere, H., Egerer, M.H. & Philpott, S.M. (2019) Environmental and spatial filtering of ladybird beetle community composition and functional traits in urban landscapes. Journal of Urban Ecology, 5: 1–12

Lister, A.M. (2011) Natural history collections as sources of long-term datasets. Trends in Ecology & Evolution, 26, 153–154.

Long, E.Y. & Finke, D.L. (2014) Contribution of predator identity to the suppression of herbivores by a diverse predator assemblage. Environmental Entomology, 43, 569–576.

Losey, J.E. & Vaughan, M. (2006) The economic value of ecological services provided by insects. BioScience, 56, 311–323.

Losey, J.E., Perlman, J.E. & Hoebeke, E.R. (2007) Citizen scientist rediscovers rare nine-spotted lady beetle, *Coccinella novemnotata*, in eastern North America. Journal of Insect Conservation, 11, 415–417.

Lövei, G.L. (1997) Global change through invasion. Nature, 388, 627–628.

Mace, G.M., Norris, K. & Fitter, A.H. (2012) Biodiversity and ecosystem services: a multilayered relationship. Trends in Ecology & Evolution, 27, 24–31.

Maes, D. & Van Dyck, H. (2001) Butterfly diversity loss in Flanders (north Belgium): Europe’s worst case scenario? Biological Conservation, 99, 263–276.

Majerus, M.E. (2016) A natural history of ladybird beetles. Cambridge University Press.

Mayer, M. & Allen, D.C. (1983) *Chilocorus stigma* (Coleoptera: Coccinellidae) and other predators of beech scale in central New York. In: Proceedings, IUFRO Beech Bark Disease Working Party Conference; 1982 September 26-October 8; Hamden, CT. USDA Forest Service, Northeastern Forest Experiment Station. Gen. Tech. Rep. WO-37. US Department of Agriculture, Forest Service: 89-98. (ed by, pp. 89–98.

Mcclure, M.S. (1986) Role of predators in regulation of endemic populations of *Matsucoccus matsumurae* (Homoptera: Margarodidae) in Japan. Environmental Entomology, 15, 976–983.

Meineke, E.K. & Daru, B.H. (2021) Bias assessments to expand research harnessing biological collections. Trends in Ecology & Evolution, 36, 1071–1082.

Meineke, E.K., Davies, T.J., Daru, B.H. & Davis, C.C. (2019) Biological collections for understanding biodiversity in the Anthropocene. Philosophical Transactions of the Royal Society B: Biological Sciences, 374, 20170386.

Michaud, J.P. (2001) Numerical response of *Olla v-nigrum* (Coleoptera: Coccinellidae) to infestations of Asian citrus psyllid, (Hemiptera: Psyllidae) in Florida. The Florida Entomologist, 84, 608–612.

Montgomery, G.A., Dunn, R.R., Fox, R., Jongejans, E., Leather, S.R., Saunders, M.E., Shortall, C.R., Tingley, M.W. & Wagner, D.L. (2020) Is the insect apocalypse upon us? How to find out. Biological Conservation, 241, 108327.

Newbold, T., Hudson, L.N., Hill, S.L.L., Contu, S., Lysenko, I., Senior, R.A., Börger, L., Bennett, D.J., Choimes, A., Collen, B., Day, J., De Palma, A., Díaz, S., Echeverria-Londoño, S., Edgar, M.J., Feldman, A., Garon, M., Harrison, M.L.K., Alhusseini, T., Ingram, D.J., Itescu, Y., Kattge, J., Kemp, V., Kirkpatrick, L., Kleyer, M., Correia, D.L.P., Martin, C.D., Meiri, S., Novosolov, M., Pan, Y., Phillips, H.R.P., Purves, D.W., Robinson, A., Simpson, J., Tuck, S.L., Weiher, E., White, H.J., Ewers, R.M., Mace, G.M., Scharlemann, J.P.W. & Purvis, A. (2015) Global effects of land use on local terrestrial biodiversity. Nature, 520, 45–50.

Oatman, E.R., Legner, E.F. & Brooks, R.F. (1964) An ecological study of arthropod populations on apple in Northeastern Wisconsin: Insect species present. Journal of Economic Entomology, 57, 978–983.

Obrycki, J.J. & Kring, T.J. (1998) Predaceous Coccinellidae in biological control. Annual Review of Entomology, 43, 295–321.

Oksanen, J., Blanchet, F.G., Kindt, R., Legendre, P., O’Hara, R.B., Simpson, G.L., Solymos, P., Stevens, M.H.H. & Wagner, H. (2011) vegan: community ecology package. R package version 2.5-6. https://CRAN.R-project.org/package=vegan.

Ortiz-Martínez, S., Staudacher, K., Baumgartner, V., Traugott, M. & Lavandero, B. (2020) Intraguild predation is independent of landscape context and does not affect the temporal dynamics of aphids in cereal fields. Journal of Pest Science, 93, 235–249.

Parker, D.M., Turo, K.J., Delgado de la flor, Y.A. & Gardiner, M.M. (2020) Landscape context influences the abundance and richness of native lady beetles occupying urban vacant land. Urban Ecosystems, 23, 1299–1310.

Pebesma, E.J. (2018) Simple features for R: Standardized support for spatial vector data. The R Journal, 10, 439.

Pell, J.K., Baverstock, J., Roy, H.E., Ware, R.L. & Majerus, M.E.N. (2008) Intraguild predation involving *Harmonia axyridis:* a review of current knowledge and future perspectives. BioControl, 53, 147–168.

Pimentel, D. & Wheeler, A.G., Jr. (1973) Species and diversity of arthropods in the alfalfa community. Environmental Entomology, 2, 659–668.

Potts, S.G., Biesmeijer, J.C., Kremen, C., Neumann, P., Schweiger, O. & Kunin, W.E. (2010) Global pollinator declines: trends, impacts and drivers. Trends in Ecology & Evolution, 25, 345–353.

Putman, W.L. (1964) Occurrence and food of some coccinellids (Coleoptera) in Ontario peach orchards. The Canadian Entomologist, 96, 1149–1155.

R Core Team (2020) R: A language and environment for statistical computing. R Foundation for Statistical Computing.

Ricci, C., Ponti, L. & Pires, A. (2005) Migratory flight and pre-diapause feeding of *Coccinella septempunctata* (Coleoptera) adults in agricultural and mountain ecosystems of Central Italy. European Journal of Entomology, 102, 531–538.

Ries, L., Zipkin, E.F. & Guralnick, R.P. (2019) Tracking trends in monarch abundance over the 20th century is currently impossible using museum records. Proceedings of the National Academy of Sciences, 116, 13745–13748.

Rocha, A.E., Souza, N.F.E., Bleakley, A.D.L., Burley, C., Mott, L.J., Rue-Glutting, G. & Fellowes, D.E.M. (2018) Influence of urbanisation and plants on the diversity and abundance of aphids and their ladybird and hoverfly predators in domestic gardens. EJE, 115, 140–149.

Rondoni, G., Borges, I., Collatz, J., Conti, E., Costamagna, A.C., Dumont, F., Evans, E.W., Grez, A.A., Howe, A.G., Lucas, E., Maisonhaute, J.-É., Onofre Soares, A., Zaviezo, T. & Cock, M.J.W. (2021) Exotic ladybirds for biological control of herbivorous insects – a review. Entomologia Experimentalis et Applicata, 169, 6–27.

Ross, N. (2019) GAMs in R: Interactive Course, https://noamross.github.io/gams-in-r-course/.

Roy, H.E., Adriaens, T., Isaac, N.J.B., Kenis, M., Onkelinx, T., Martin, G.S., Brown, P.M.J., Hautier, L., Poland, R., Roy, D.B., Comont, R., Eschen, R., Frost, R., Zindel, R., Van Vlaenderen, J., Nedvěd, O., Ravn, H.P., Grégoire, J.-C., de Biseau, J.-C. & Maes, D. (2012) Invasive alien predator causes rapid declines of native European ladybirds. Diversity and Distributions, 18, 717–725.

Roy, H.E., Brown, P.M.J., Adriaens, T., Berkvens, N., Borges, I., Clusella-Trullas, S., Comont, R.F., De Clercq, P., Eschen, R., Estoup, A., Evans, E.W., Facon, B., Gardiner, M.M., Gil, A., Grez, A.A., Guillemaud, T., Haelewaters, D., Herz, A., Honek, A., Howe, A.G., Hui, C., Hutchison, W.D., Kenis, M., Koch, R.L., Kulfan, J., Lawson Handley, L., Lombaert, E., Loomans, A., Losey, J., Lukashuk, A.O., Maes, D., Magro, A., Murray, K.M., Martin, G.S., Martinkova, Z., Minnaar, I.A., Nedved, O., Orlova-Bienkowskaja, M.J., Osawa, N., Rabitsch, W., Ravn, H.P., Rondoni, G., Rorke, S.L., Ryndevich, S.K., Saethre, M.-G., Sloggett, J.J., Soares, A.O., Stals, R., Tinsley, M.C., Vandereycken, A., van Wielink, P., Viglášová, S., Zach, P., Zakharov, I.A., Zaviezo, T. & Zhao, Z. (2016) The harlequin ladybird, *Harmonia axyridis*: Global perspectives on invasion history and ecology. Biological Invasions, 18, 997–1044.

Rusch, A., Chaplin-Kramer, R., Gardiner, M.M., Hawro, V., Holland, J., Landis, D., Thies, C., Tscharntke, T., Weisser, W.W., Winqvist, C., Woltz, M. & Bommarco, R. (2016) Agricultural landscape simplification reduces natural pest control: A quantitative synthesis. Agriculture, Ecosystems & Environment, 221, 198–204.

Sánchez-Bayo, F. & Wyckhuys, K.A.G. (2019) Worldwide decline of the entomofauna: A review of its drivers. Biological Conservation, 232, 8–27.

Saunders, M.E., Janes, J.K. & O’Hanlon, J.C. (2020) Moving on from the insect apocalypse narrative: Engaging with evidence-based insect conservation. BioScience, 70, 80–89.

Schowalter, T.D., Pandey, M., Presley, S.J., Willig, M.R. & Zimmerman, J.K. (2021) Arthropods are not declining but are responsive to disturbance in the Luquillo Experimental Forest, Puerto Rico. Proceedings of the National Academy of Sciences, 118, e2002556117.

Schuch, S., Wesche, K. & Schaefer, M. (2012) Long-term decline in the abundance of leafhoppers and planthoppers (Auchenorrhyncha) in Central European protected dry grasslands. Biological Conservation, 149, 75–83.

Seibold, S., Gossner, M.M., Simons, N.K., Blüthgen, N., Müller, J., Ambarlı, D., Ammer, C., Bauhus, J., Fischer, M., Habel, J.C., Linsenmair, K.E., Nauss, T., Penone, C., Prati, D., Schall, P., Schulze, E.-D., Vogt, J., Wöllauer, S. & Weisser, W.W. (2019) Arthropod decline in grasslands and forests is associated with landscape-level drivers. Nature, 574, 671–674.

Shaffer, H.B., Fisher, R.N. & Davidson, C. (1998) The role of natural history collections in documenting species declines. Trends in Ecology & Evolution, 13 1, 27–30.

Shortall, C.R., Moore, A., Smith, E., Hall, M.J., Woiwod, I.P. & Harrington, R. (2009) Long-term changes in the abundance of flying insects. Insect Conservation and Diversity, 2, 251–260.

Simmons, B.I., Balmford, A., Bladon, A.J., Christie, A.P., De Palma, A., Dicks, L.V., Gallego-Zamorano, J., Johnston, A., Martin, P.A., Purvis, A., Rocha, R., Wauchope, H.S., Wordley, C.F.R., Worthington, T.A. & Finch, T. (2019) Worldwide insect declines: An important message, but interpret with caution. Ecology and Evolution, 9, 3678–3680.

Sloggett, J.J. (2017) *Harmonia axyridis* (Coleoptera: Coccinellidae): Smelling the rat in native ladybird declines. European Journal of Entomology, 114, 455–461.

Smith, B.C. (1971) Effects of various factors on the local distribution and density of coccinellid adults on corn (Coleoptera: Coccinellidae). The Canadian Entomologist, 103, 1115–1120.

Smith, C.A. & Gardiner, M.M. (2013) Biodiversity loss following the introduction of exotic competitors: Does intraguild predation explain the decline of native lady beetles? PLoS ONE, 8, e84448.

Snyder, W.E. (2009) Coccinellids in diverse communities: Which niche fits? Biological Control, 51, 323–335.

Snyder, W.E. & Evans, E.W. (2006) Ecological effects of invasive arthropod generalist predators. Annu. Rev. Ecol. Evol. Syst., 37, 95–122.

Snyder, W.E., Clevenger, G.M. & Eigenbrode, S.D. (2004) Intraguild predation and successful invasion by introduced ladybird beetles. Oecologia, 140, 559–565.

Sohl, T., Reker, R., Bouchard, M., Sayler, K., Dornbierer, J., Wika, S., Quenzer, R. & Friesz, A. (2016) Modeled historical land use and land cover for the conterminous United States. Journal of Land Use Science, 11, 476–499.

Sohl, T., Reker, R., Bouchard, M., Sayler, K., Dornbierer, J., Wika, S., Quenzer, R. & Friesz, A. (2018) Modeled historical land use and land cover for the conterminous United States: 1938-1992: U.S. Geological Survey data release. https://doi.org/10.5066/F7KK99RR.

Staines, C.L. (2008) Coccinellidae or ladybird beetles (Insecta: Coleoptera) of Plummers Island, Maryland. Bulletin of the Biological Society of Washington, 15, 149–150.

Staines Jr, C., Rothschild, M. & Trumbule, R. (1990) A survey of the Coccinellidae (Coleoptera) associated with nursery stock in Maryland. Proceedings of the Entomological Society of Washington, 92, 310–313.

Steffens, W.P. & Lumen, R. (2015) Decline in relative abundance of *Hippodamia convergens* (Coleoptera: Coccinellidae) in fall shoreline aggregations on western Lake Superior. Great Lakes Entomologist, 48, 8.

Suarez, A.V. & Tsutsui, N.D. (2004) The value of museum collections for research and society. BioScience, 54, 66–74.

Thomas, A.P., Trotman, J., Wheatley, A., Aebi, A., Zindel, R. & Brown, P.M.J. (2013) Predation of native coccinellids by the invasive alien *Harmonia axyridis* (Coleoptera: Coccinellidae): detection in Britain by PCR-based gut analysis. Insect Conservation and Diversity, 6, 20–27.

Thomas, C.D., Jones, T.H. & Hartley, S.E. (2019) “Insectageddon”: A call for more robust data and rigorous analyses. Global Change Biology, 25, 1891–1892.

Turnock, W.J., Wise, I.L. & Matheson, F.O. (2003) Abundance of some native coccinellines (Coleoptera: Coccinellidae) before and after the appearance of Coccinella septempunctata. The Canadian Entomologist, 135, 391–404.

Vitousek, P.M., Mooney, H.A., Lubchenco, J. & Melillo, J.M. (1997) Human domination of Earth’s ecosystems. Science, 277, 494–499.

Wagner, D.L. (2020) Insect declines in the anthropocene. Annual Review of Entomology, 65, 457–480.

Wagner, D.L., Grames, E.M., Forister, M.L., Berenbaum, M.R. & Stopak, D. (2021) Insect decline in the Anthropocene: Death by a thousand cuts. Proceedings of the National Academy of Sciences, 118, e2023989118.

Warren, M.S., Maes, D., van Swaay, C.A.M., Goffart, P., Van Dyck, H., Bourn, N.A.D., Wynhoff, I., Hoare, D. & Ellis, S. (2021) The decline of butterflies in Europe: Problems, significance, and possible solutions. Proceedings of the National Academy of Sciences, 118, e2002551117.

Weber, D.C. & Lundgren, J.G. (2009) Assessing the trophic ecology of the Coccinellidae: Their roles as predators and as prey. Biological Control, 51, 199–214.

Wheeler Jr, A. & Hoebeke, E. (1995) *Coccinella novemnotata* in northeastern North America: Historical occurrence and current status (Coleoptera: Coccinellidae). Entomological Society of Washington (USA), 97(3): pp.701–716.

Wickham, J., Stehman, S.V., Sorenson, D.G., Gass, L. & Dewitz, J.A. (2021) Thematic accuracy assessment of the NLCD 2016 land cover for the conterminous United States. Remote Sensing of Environment, 257, 112357.

Wilcove, D.S., Rothstein, D., Jason, D., Phillips, A. & Losos, E. (1998) Quantifying threats to imperiled species in the United States. BioScience, 48, 607–615.

Winker, K. (2004) Natural history museums in a postbiodiversity era. BioScience, 54, 455–459.

Woltz, M.J. & Landis, D.A. (2014) Coccinellid response to landscape composition and configuration. Agricultural and Forest Entomology, 16, 341–349.

Wood, S.N. (2017) Generalized Additive Models: An Introduction with R (2nd edition). Chapman and Hall/CRC.

Yang, L.H. & Gratton, C. (2014) Insects as drivers of ecosystem processes. Current Opinion in Insect Science, 2, 26–32.

Zaviezo, T., Grez, A.A., Estades, C.F. & Perez, A. (2006) Effects of habitat loss, habitat fragmentation, and isolation on the density, species richness, and distribution of ladybeetles in manipulated alfalfa landscapes. Ecological Entomology, 31, 646–656.

