## Appendix 1 for "Landscape change and alien invasions drive shifts in native lady beetle communities over a century"

**Table S1.1.** Ohio lady beetle records were compiled from the 25 institutions listed below with assistance from their curators.

| Institution | Curators and/or Contacts |
|---|---|
| California Academy of Sciences | Susan Gin and Christopher C. Grinter |
| Natural History Museum of Los Angeles County | Brian Brown and Weiping Xie |
| Peabody Museum, Yale University | Lawrence Gall |
| Biodiversity Research Collections, University of Connecticut | Jane O'Donnell |
| Smithsonian National Museum of Natural History | Natalie Vandenburgh |
| Collection of Arthropods, University of Georgia | Joseph V. McHugh and E. Richard Hoebeke |
| Gantz Family Collections Center, The Field Museum | Rebekah Shuman Baquiran |
| Illinois Natural History Survey, University of Illinois | Thomas McElrath |
| Louisiana State Arthropod Museum, Louisiana State University | Victoria Moseley Bayless |
| A.J. Cook Arthropod Research Collection, Michigan State University | Gary L. Parsons |
| Museum of Zoology, University of Michigan | Erika Tucker |
| University of Minnesota | Robin Thomson |
| Wilbur R. Enns Entomology Museum, University of Missouri | Kristin B. Simpson |
| Mississippi Entomological Museum, Mississippi State University | Terence L. Schiefer |
| C.A. Triplehorn Insect Collection, The Ohio State University | Luciana Musetti |
| Agricultural Technical Institute Teaching Collection, The Ohio State University | Jon Van Gray |
| Cleveland Museum of Natural History | Nicole Gunter |
| Carnegie Museum of Natural History | Robert Androw and Robert Davidson |
| The Frost Entomological Museum, Pennsylvania State University | Online SCAN Project Database (https://scan-bugs.org/portal/collections/misc/collprofiles.php?collid=126) |
| Clemson University Arthropod Collection, Clemson University | Michael Ferro |
| Severin-McDaniel Insect Research Collection, South Dakota State University | Louis Hesler |
| Brigham Young University | Shawn M. Clark |
| Crane Hollow Nature Preserve | Gary and Holly Coovert |
| Boonshoft Museum of Discovery | Kaitlin U. Campbell and Thomas O. Crist |
| The Lost Ladybug Project (http://www.lostladybug.org/) | Rebecca Smyth |
