## Appendix 2 for "Landscape change and alien invasions drive shifts in native lady beetle communities over a century"

**Table S2.1.** NLCD reclassification crosswalk table. Version NLCD 2016 was used in the analysis (Dewitz 2019).

| **Map Class** | **NLCD Classification Description (Class Value)** |
| --- | --- |
| Agriculture | Hay/Pasture (81)<br>Cultivated Crops (82) |
| Developed | Developed, Open Space (21)<br>Developed, Low Intensity (22)<br>Developed, Medium Intensity (23)<br>Developed, High Intensity (24) |
| Forest | Deciduous Forest (41)<br>Evergreen Forest (42)<br>Mixed Forest (43)<br>Woody Wetlands (90) |
| Non-target | Open Water (11)<br>Perennial Snow/Ice (12)<br>Barren Land (31)<br>Shrub/Scrub (52)<br>Herbaceous (71)<br>Emergent Herbaceous Wetlands (95) |

Dewitz, J. (2019) National Land Cover Database (NLCD) 2016 Products (ver. 2.0, July 2020): U.S. Geological Survey data release. https://doi.org/10.5066/P96HHBIE.

**Table S2.2.** Historical LULC backcast reclassification crosswalk table for years 1938, 1970, 1992. Version from Sohl et al. (2018) was used in the analysis.

| **Map Class** | **Backcast Classification Description (Class Value)** |
|---|---|
| Agriculture | Cultivated Cropland (13)<br>Hay/Pasture (14) |
| Developed | Urban/Developed (2)<br>Mining (6) |
| Forest | Deciduous Forest (8)<br>Evergreen Forest (9)<br>Mixed Forest (10)<br>Woody Wetland (16) |
| Non-target | Open Water (1)<br>Barren (7)<br>Grassland (11)<br>Shrubland (12)<br>Herbaceous Wetland (15) |

Sohl, T., Reker, R., Bouchard, M., Sayler, K., Dornbierer, J., Wika, S., Quenzer, R. & Friesz, A. (2018) Modeled historical land use and land cover for the conterminous United States: 1938-1992: U.S. Geological Survey data release. https://doi.org/10.5066/F7KK99RR.

**Table S2.3.** Ohio lady beetle records from institutions across the United States (1900-2018). We compiled historic data records from 25 institutions across the US and identified 4,194 specimens from 28 lady beetle species. Data collection was focused on Coccinellinae specimens and four additional species: *Brachiacantha ursina*, *Chilocorus stigma*, *Hyperaspis undulata*, and *Psyllobora vigintimaculata*.

| **Lady beetle species** | **1900** | **1910** | **1920** | **1930** | **1940** | **1950** | **1960** | **1970** | **1980** | **1990** | **2000** | **2010** | **Total** |
|---|---|---|---|---|---|---|---|---|---|---|---|---|---|
| *Adalia bipunctata* | 6 | 8 | 8 | 49 | 18 | 22 | 66 | 15 | 20 | 6 | 0 | 0 | 218 |
| *Anatis labiculata* | 1 | 10 | 4 | 26 | 36 | 23 | 27 | 6 | 18 | 19 | 1 | 5 | 176 |
| *Anatis mali* | 0 | 0 | 0 | 5 | 1 | 5 | 4 | 3 | 1 | 0 | 2 | 2 | 23 |
| *Anisosticta bitriangularis* | 0 | 0 | 5 | 0 | 0 | 0 | 0 | 1 | 0 | 0 | 0 | 10 | 16 |
| *Brachiacantha ursina* | 8 | 19 | 19 | 111 | 20 | 27 | 21 | 6 | 5 | 3 | 12 | 4 | 255 |
| *Chilocorus stigma* | 18 | 10 | 6 | 12 | 5 | 4 | 17 | 2 | 13 | 13 | 7 | 4 | 111 |
| *Coccinella novemnotata* | 10 | 21 | 28 | 41 | 7 | 28 | 16 | 13 | 5 | 0 | 0 | 0 | 169 |
| *Coccinella septempunctata* | 0 | 0 | 0 | 0 | 0 | 0 | 0 | 4 | 47 | 6 | 19 | 26 | 102 |
| *Coccinella transversoguttata* | 0 | 0 | 5 | 4 | 0 | 4 | 3 | 25 | 10 | 0 | 0 | 0 | 51 |
| *Coccinella trifasciata* | 0 | 3 | 5 | 4 | 0 | 1 | 5 | 1 | 0 | 0 | 0 | 0 | 19 |
| *Coccinella undecimpunctata* | 0 | 0 | 0 | 0 | 0 | 1 | 0 | 0 | 0 | 0 | 0 | 0 | 1 |
| *Coleomegilla maculata* | 11 | 14 | 32 | 135 | 11 | 22 | 71 | 55 | 263 | 28 | 9 | 39 | 690 |
| *Cycloneda munda* | 10 | 4 | 13 | 28 | 10 | 16 | 46 | 32 | 65 | 10 | 48 | 20 | 302 |
| *Harmonia axyridis* | 0 | 0 | 0 | 0 | 0 | 0 | 0 | 0 | 0 | 35 | 199 | 236 | 470 |
| *Hippodamia convergens* | 6 | 14 | 15 | 131 | 23 | 30 | 66 | 37 | 62 | 1 | 1 | 14 | 400 |
| *Hippodamia glacialis* | 1 | 4 | 3 | 13 | 4 | 1 | 1 | 5 | 2 | 0 | 3 | 2 | 39 |
| *Hippodamia parenthesis* | 20 | 12 | 51 | 102 | 22 | 20 | 30 | 29 | 40 | 0 | 13 | 14 | 353 |
| *Hippodamia quindecimmaculata* | 2 | 0 | 0 | 1 | 0 | 0 | 0 | 0 | 0 | 0 | 0 | 0 | 3 |
| *Hippodamia tredecimpunctata* | 6 | 5 | 8 | 82 | 6 | 4 | 31 | 5 | 5 | 0 | 0 | 3 | 155 |
| *Hippodamia variegata* | 0 | 0 | 0 | 0 | 0 | 0 | 0 | 0 | 0 | 0 | 4 | 6 | 10 |
| *Hyperaspis undulata* | 5 | 3 | 6 | 28 | 1 | 3 | 5 | 2 | 3 | 2 | 3 | 0 | 61 |
| *Mulsantina luteodorsa* | 0 | 0 | 0 | 0 | 0 | 0 | 0 | 0 | 0 | 0 | 1 | 0 | 1 |
| *Mulsantina picta* | 0 | 0 | 2 | 16 | 12 | 3 | 11 | 3 | 1 | 0 | 2 | 1 | 51 |

| | | | | | | | | | | | | | |
|---|---|---|---|---|---|---|---|---|---|---|---|---|---|
| *Myzia pullata* | 0 | 0 | 0 | 12 | 5 | 6 | 1 | 1 | 1 | 0 | 3 | 1 | 30 |
| *Neoharmonia venusta* | 2 | 0 | 0 | 63 | 32 | 13 | 3 | 1 | 2 | 1 | 1 | 1 | 119 |
| *Olla v-nigrum* | 0 | 0 | 0 | 5 | 0 | 14 | 0 | 0 | 1 | 0 | 1 | 0 | 21 |
| *Propylea quatuordecimpunctata* | 0 | 0 | 0 | 0 | 0 | 0 | 0 | 0 | 0 | 0 | 6 | 21 | 27 |
| *Psyllobora vigintimaculata* | 2 | 5 | 8 | 95 | 11 | 11 | 9 | 3 | 51 | 60 | 61 | 5 | 321 |
| **Total Collections** | **108** | **132** | **218** | **963** | **224** | **258** | **433** | **249** | **615** | **184** | **396** | **414** | **4194** |
| **Total Collections of Natives** | **108** | **132** | **218** | **963** | **224** | **257** | **433** | **245** | **568** | **143** | **168** | **125** | **3584** |
| **Total Species** | **15** | **14** | **17** | **21** | **17** | **21** | **19** | **21** | **20** | **12** | **20** | **19** | **28** |
| **Total Native Species** | **15** | **14** | **17** | **21** | **17** | **20** | **19** | **20** | **19** | **10** | **16** | **15** | **23** |

**Table S2.4.** Results for permutational multivariate analysis of variance (PERMANOVA) and analysis of multivariate homogeneity of group dispersions (BETADISPER) for taxonomic community similarity of all lady beetle species and aphidophagous lady beetle species across decades in Ohio, USA. PERMANOVA tests whether the centroid of communities differs among groups in multivariate space, while BETADISPER tests whether groups differ in the amount of dispersion from its spatial median among communities within a group. Beta-diversity indices were incidence-based pairwise Sorensen dissimilarity ($\beta_{sor}$) which was partitioned into turnover ($\beta_{sim}$; reflects species replacement) and nestedness ($\beta_{sne}$; reflects species loss or gain) components. Significant differences ($\alpha < 0.05$) in lady beetle community similarity are indicated in bold.

| Decades | Test | | All Lady Beetle Species | | | Aphidophagous Species | | |
|---|---|---|---|---|---|---|---|---|
| | | | ($\beta_{sim}$) | ($\beta_{sne}$) | ($\beta_{sor}$) | ($\beta_{sim}$) | ($\beta_{sne}$) | ($\beta_{sor}$) |
| 1900s-1910s | PERMANOVA | *F* | 4.56 | -1.04 | 2.35 | 4.35 | -1.08 | 2.23 |
| | | *P* | 0.111 | 0.923 | 0.066 | 0.155 | 0.963 | 0.102 |
| | BETADISPER | *F* | 5.10 | 2.78 | 0.09 | 6.23 | 1.12 | 0.06 |
| | | *P* | **0.043** | 0.121 | 0.760 | **0.028** | 0.310 | 0.810 |
| 1910s-1920s | PERMANOVA | *F* | 0.53 | 24.92 | 1.01 | 0.08 | 49.49 | 0.60 |
| | | *P* | 0.571 | **0.011** | 0.466 | 0.657 | **0.003** | 0.633 |
| | BETADISPER | *F* | 4.75 | 1.99 | 0.99 | 2.27 | 0.06 | 2.37 |
| | | *P* | **0.049** | 0.183 | 0.337 | 0.157 | 0.805 | 0.149 |
| 1920s-1930s | PERMANOVA | *F* | 2.19 | 5.51 | 3.88 | 2.57 | 3.38 | 2.99 |
| | | *P* | 0.301 | 0.060 | **0.014** | 0.241 | 0.120 | **0.025** |
| | BETADISPER | *F* | 0.93 | 0.49 | 1.45 | 1.26 | 0.32 | 2.29 |
| | | *P* | 0.352 | 0.495 | 0.250 | 0.282 | 0.580 | 0.155 |
| 1930s-1940s | PERMANOVA | *F* | -1.73 | 5.86 | 3.12 | -68.35 | 5.68 | 2.96 |
| | | *P* | 0.866 | 0.073 | **0.003** | 0.997 | 0.056 | **0.008** |
| | BETADISPER | *F* | 4.75 | 1.16 | 5.80 | 3.91 | 1.85 | 5.45 |
| | | *P* | **0.046** | 0.299 | **0.030** | 0.067 | 0.194 | **0.034** |
| 1940s-1950s | PERMANOVA | *F* | 0.33 | 2.08 | 1.06 | -3.44 | 2.70 | 1.19 |
| | | *P* | 0.614 | 0.230 | 0.376 | 0.967 | 0.159 | 0.381 |
| | BETADISPER | *F* | 0.56 | 0.28 | 0.33 | 0.25 | 1.19 | 0.24 |
| | | *P* | 0.465 | 0.604 | 0.572 | 0.619 | 0.292 | 0.626 |
| 1950s-1960s | PERMANOVA | *F* | 2.23 | 6.65 | 1.10 | 1.73 | 13.04 | 0.91 |

| | | | | | | | | |
|---|---|---|---|---|---|---|---|---|
| | | *P* | 0.215 | **0.037** | 0.387 | 0.313 | **0.004** | 0.483 |
| | BETADISPER | *F* | 0.002 | 0.58 | 0.39 | 0.01 | 2.55 | 1.31 |
| | | *P* | 0.963 | 0.456 | 0.539 | 0.917 | 0.132 | 0.270 |
| 1960s-1970s | PERMANOVA | *F* | 0.05 | 1.63 | 0.46 | 1.38 | 0.45 | 0.92 |
| | | *P* | 0.857 | 0.277 | 0.805 | 0.348 | 0.533 | 0.442 |
| | BETADISPER | *F* | 3.25 | 5.69 | 0.92 | 1.10 | 0.30 | 2.06 |
| | | *P* | 0.092 | **0.031** | 0.352 | 0.311 | 0.591 | 0.172 |
| 1970s-1980s | PERMANOVA | *F* | 0.03 | -0.14 | 0.33 | 0.60 | 0.55 | 0.43 |
| | | *P* | 0.717 | 0.735 | 0.947 | 0.636 | 0.483 | 0.812 |
| | BETADISPER | *F* | 1.12 | 2.26 | 0.01 | 1.57 | 0.58 | 1.06 |
| | | *P* | 0.305 | 0.154 | 0.943 | 0.231 | 0.459 | 0.320 |
| 1980s-1990s | PERMANOVA | *F* | 4.32 | 0.50 | 2.94 | 1.73 | 32.80 | 4.73 |
| | | *P* | 0.091 | 0.570 | **0.021** | 0.260 | **0.004** | **0.012** |
| | BETADISPER | *F* | 0.42 | 0.32 | 0.18 | 1.62 | 0.39 | 1.89 |
| | | *P* | 0.527 | 0.577 | 0.672 | 0.226 | 0.540 | 0.194 |
| 1990s-2000s | PERMANOVA | *F* | 4.98 | 0.22 | 3.36 | 3.32 | 2.80 | 4.00 |
| | | *P* | 0.061 | 0.670 | **0.009** | 0.151 | 0.171 | **0.010** |
| | BETADISPER | *F* | 2.73 | 0.47 | 0.53 | 3.52 | 0.99 | 0.33 |
| | | *P* | 0.122 | 0.504 | 0.479 | 0.083 | 0.335 | 0.570 |
| 2000s-2010s | PERMANOVA | *F* | 2.24 | 0.08 | 1.58 | 0.24 | 2.49 | 0.75 |
| | | *P* | 0.190 | 0.960 | 0.187 | 0.709 | 0.108 | 0.605 |
| | BETADISPER | *F* | 0.04 | 0.47 | 0.25 | 0.001 | 0.40 | 0.56 |
| | | *P* | 0.832 | 0.501 | 0.624 | 0.994 | 0.536 | 0.465 |

**Figure S2.1.** Number of collections of alien lady beetle species in Ohio by decade from 1900 to 2018. All alien species are aphidophagous.

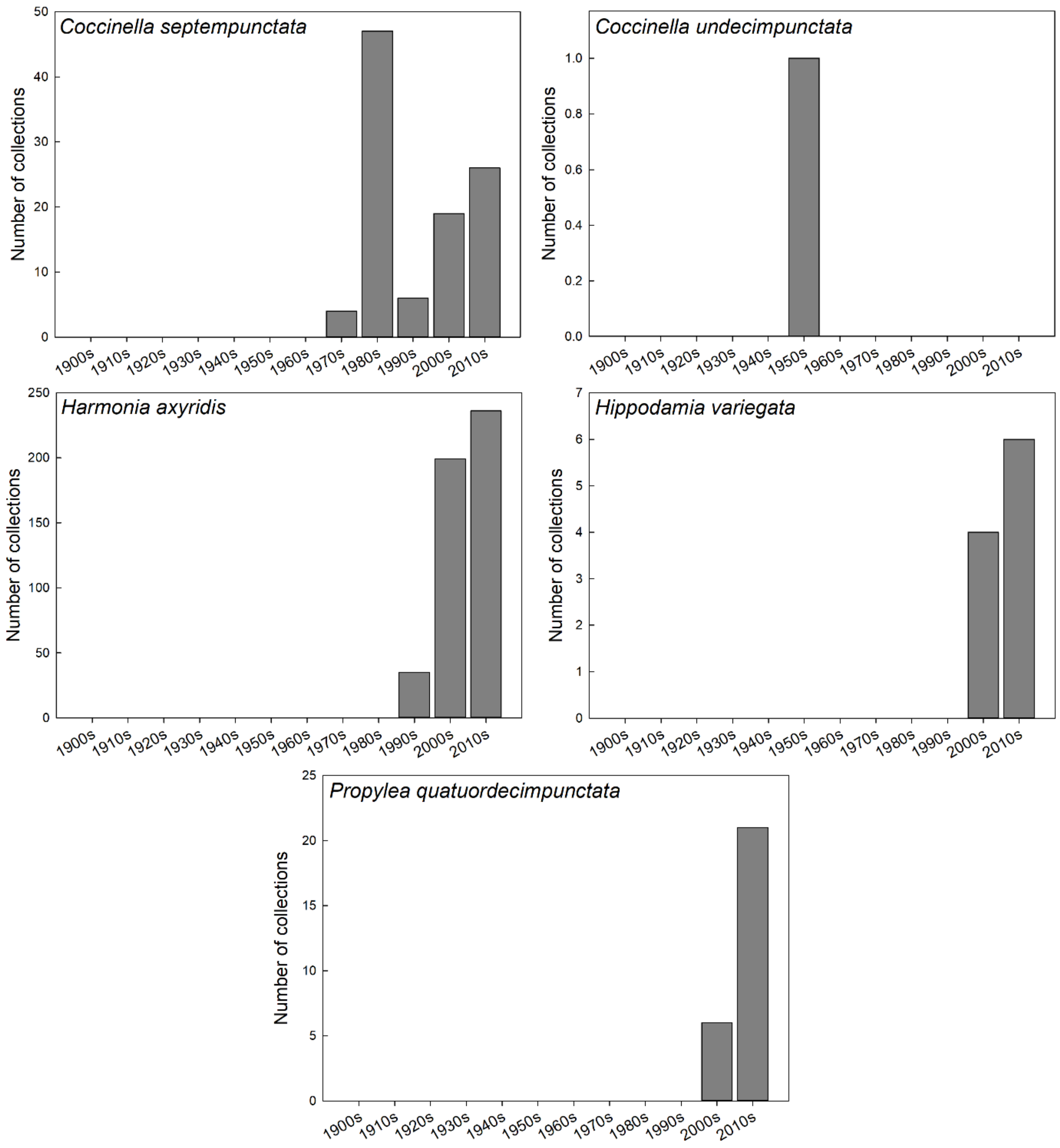

**Figure S2.2.** Number of collections of native aphidophagous lady beetle species in Ohio by decade from 1900 to 2018.

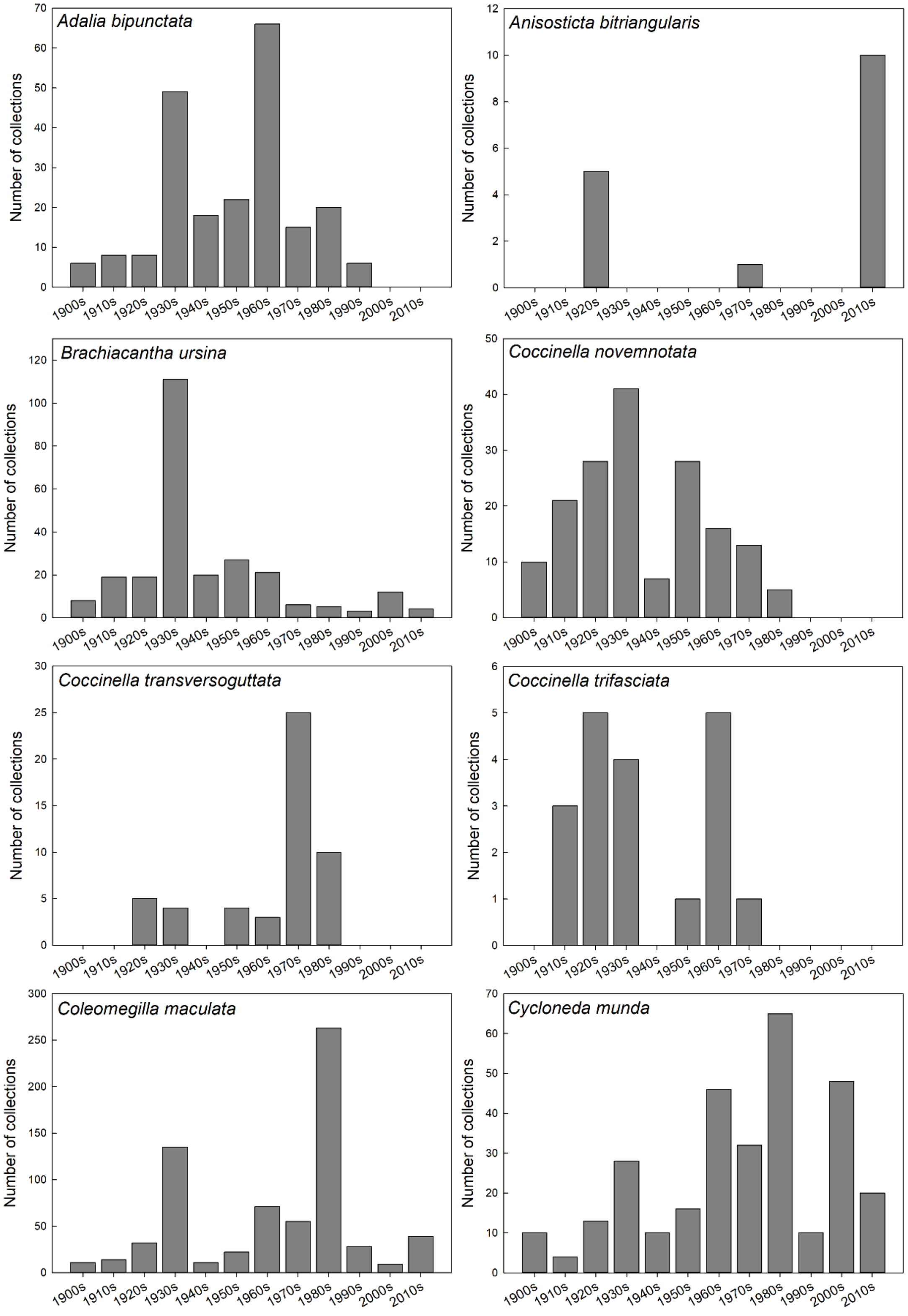

**Figure S2.3.** Number of collections of native aphidophagous lady beetle species (continued) in Ohio by decade from 1900 to 2018.

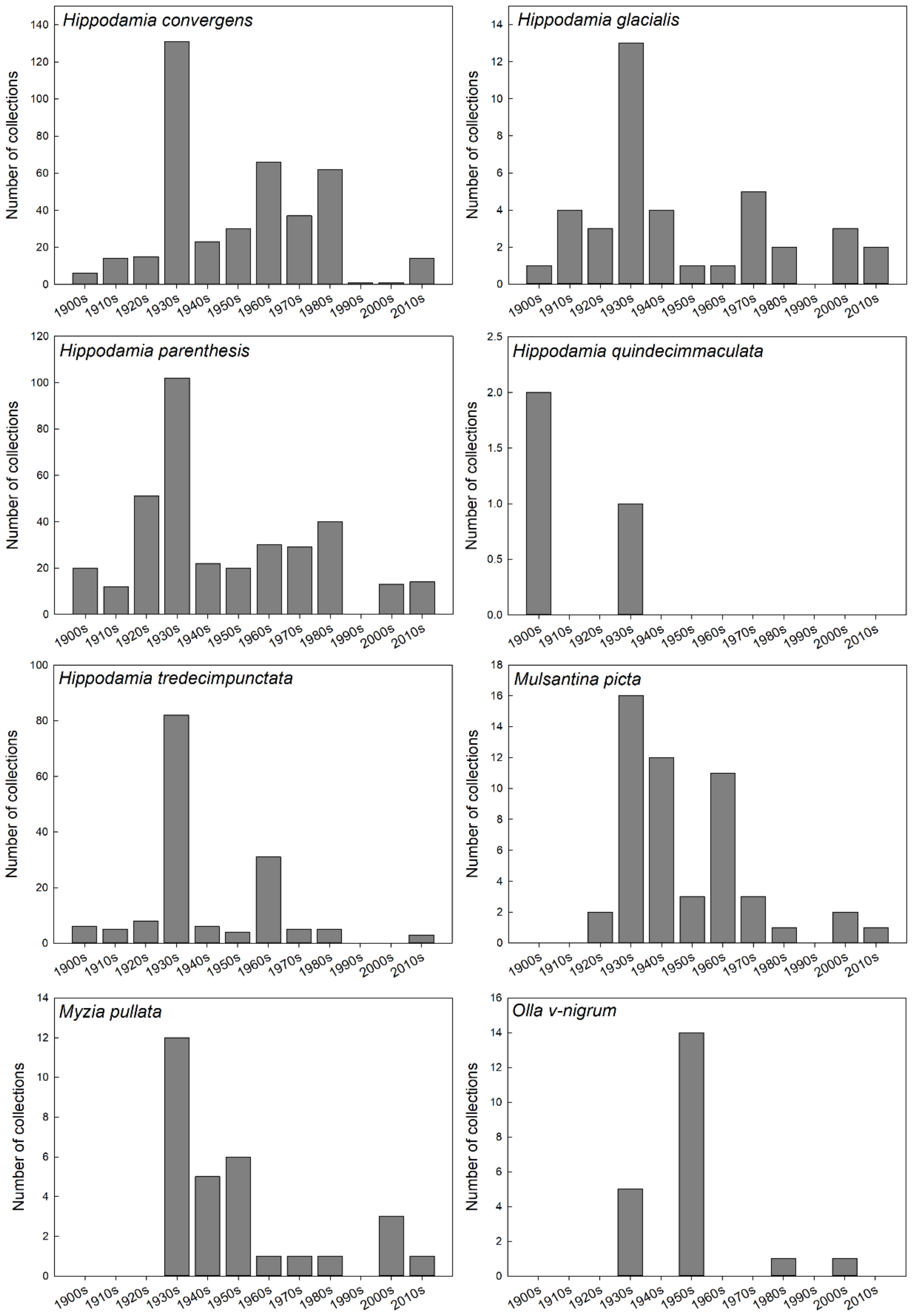

**Figure S2.4.** Number of collections of native coccidophagous and fungivorus lady beetle species in Ohio by decade from 1900 to 2018.

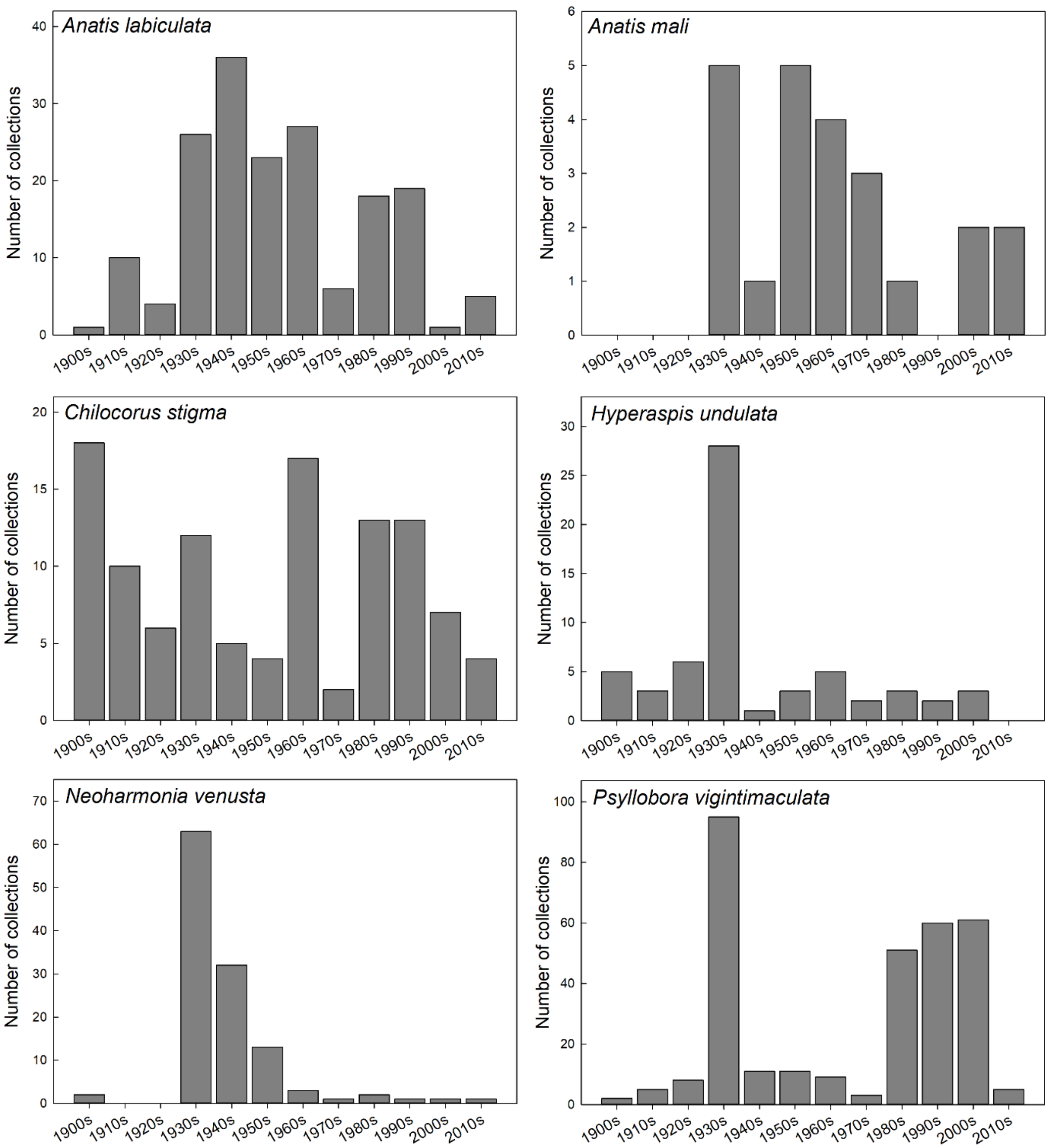

**Figure S2.5.** Non-metric multidimensional scaling (NMDS) ordination depicting incidence-based pairwise Sorensen dissimilarity ($\beta_{sor}$) for lady beetle museum records across decades in Ohio, USA. Results for permutational multivariate analysis of variance (PERMANOVA) are provided in Appendix 1: Table 2 and significant comparisons between (A) the 1920s and 1930s; (B) the 1930s and 1940s; (C) the 1980s and 1990s; and (D) the 1990s and 2000s are depicted here.

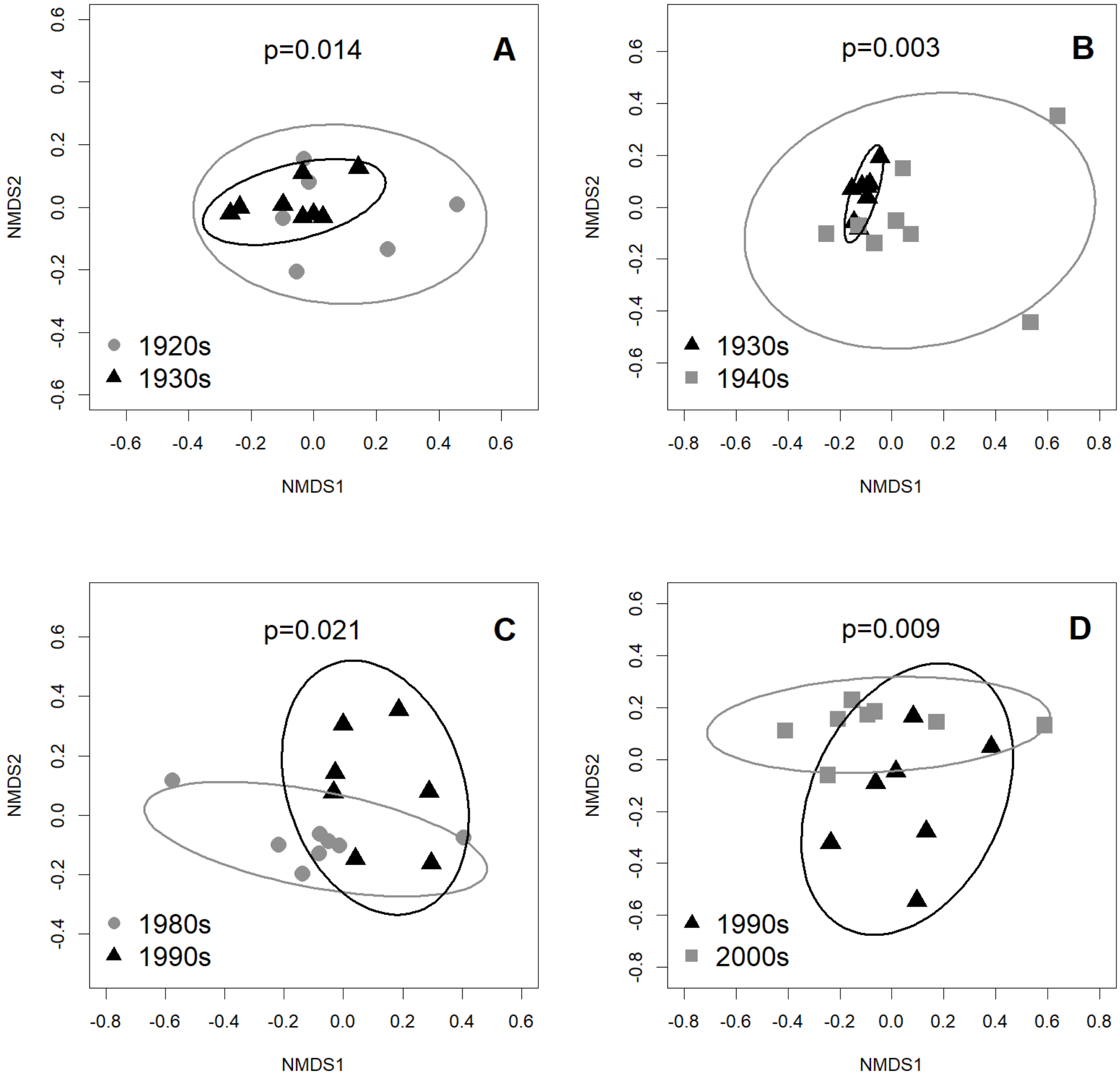

**Figure S2.6.** The variability in Ohio land cover area by county. Horizontal black lines represent the median. Boxes extend to the 25th and 75th percentile. Whiskers extend to the most extreme data point, which is no more than 1.5 times the interquartile range from the box. Data beyond the end of the whiskers are outlying points and are plotted individually.

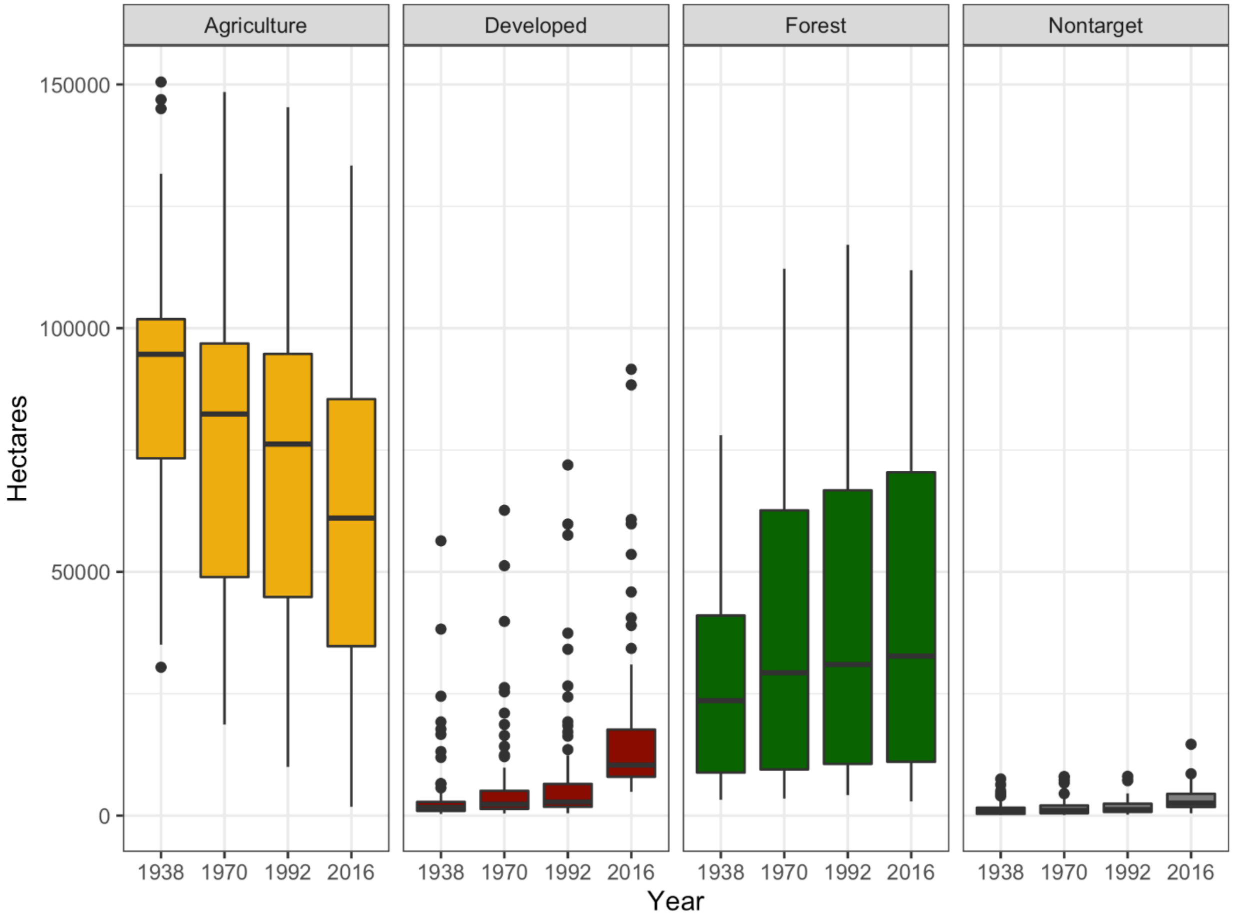
