## Appendix 3 for "Landscape change and alien invasions drive shifts in native lady beetle communities over a century"

Results from model selection and spatial analysis of responses for five key native species to spatiotemporal, invasion, and landscape parameters. Negative binomial generalized additive models used Ohio museum collections, 1930-2018 data on lady beetle captures, compiled by county and decade of capture, and were adjusted for sampling effort.

**Table S3.1.** Model selection and fit statistics for selected lady beetle species.

| Target Species | Spatial | Decade | Decade | Longitude | Latitude | Landscape metrics | Total invasive | Invasive | C7 | Harmonia | Adj R2 | Deviance explained % | -REML | AIC | Final model | Notes |
|---|---|---|---|---|---|---|---|---|---|---|---|---|---|---|---|---|
| *Coleomegilla maculata* | X | | | | | | | | | | 0.391 | 10.9 | 807.7 | 1608.2 | | |
| | X | X | | | | | | | | | 0.381 | 21.9 | 795.63 | 1585.6 | | |
| | | | X | X | X | Agriculture, Forest, Developed | X | | | | 0.4848 | 15.3 | 801.94 | 1590.28 | | |
| | | | X | X | X | Agriculture, Forest, Developed | | X | | | 0.399 | 21.4 | 782.7 | 1550.434 | | |
| | | | X | X | X | Agriculture, Forest, Developed | | X | X | X | 0.494 | 21.7 | 783.98 | 1552.709 | | |
| | | | X | | X | Agriculture, Forest, Developed | | X | X | X | 0.479 | 21.3 | 781.47 | 1548.45 | | |
| | | | X | | X | Agriculture, Forest | | X | X | X | 0.479 | 20.9 | 781.79 | 1546.83 | | |
| | | | X | | X | Agriculture | | X | X | X | 0.479 | 20.6 | 780.53 | 1543.88 | | |
| | | | X | | X | Agriculture | | X | | X | 0.483 | 20.5 | 779.7 | 1542.24 | | |
| | | | X | | X | Agriculture | | X | | | 0.404 | 20.4 | 778.87 | 1541.03 | X | |
| | | | X | | X | | | X | | | 0.301 | 19.1 | 782.55 | 1551.126 | | |

| | | | | | | | | | | | | | | | | |
|---|---|---|---|---|---|---|---|---|---|---|---|---|---|---|---|---|
| *Coccinella novemnota* | X | | | | | | | | | | 0.301 | 1.85 | 221.25 | 445.4 | | |
| | X | X | | | | | | | | | 0.328 | 7.92 | 217.61 | 447.3 | | |
| | | | X | X | X | Agriculture, Forest, Developed | X | | | | -0.363 | 31.1 | 197.62 | 400.2 | | Total invasive needs to be limited to k=4 |
| | | | X | X | X | Agriculture, Forest, Developed | | X | | | -0.347 | 31.3 | 196.74 | 400.4 | | |
| | | | X | X | X | Agriculture, Forest, Developed | | X | X | NA | -0.358 | 31.2 | 197.54 | 402.72 | | |
| | | | X | X | X | Agriculture, Forest, Developed | | | | | -0.384 | 31 | 197.04 | 398.44 | | |
| | | | X | X | X | Agriculture, Developed | | | | | -0.234 | 30.2 | 199.69 | 396.9 | | |
| | | | X | X | X | Agriculture | | | | | 0.01177 | 28.9 | 199.73 | 396.5 | X | |
| | | | X | X | | Agriculture | | | | | 0.302 | 23.3 | 202.51 | 404.25 | | |
| *Adalia bipunctata* | X | | | | | | | | | | 0.46 | 10.5 | 328.2 | 655.9 | | |
| | X | X | | | | | | | | | 0.538 | 21 | 316.39 | 638.8 | | |
| | | | X | X | X | Agriculture, Forest, Developed | X | | | | 0.436 | 19 | 312.32 | 624.71 | | Total invasive needs to be limited to k=4 |
| | | | X | X | X | Agriculture, Forest, Developed | | X | | | 0.398 | 19.9 | 308.67 | 621.6 | | |
| | | | X | X | X | Agriculture, Forest, Developed | | X | X | NA | 0.423 | 20.2 | 309.69 | 622.8 | | |
| | | | X | X | X | Agriculture, Developed | | X | | | 0.388 | 19.8 | 309.81 | 618.88 | | |
| | | | | X | X | Agriculture, Developed | | X | | | 0.324 | 19.1 | 308.8 | 617.29 | | |
| | | | | X | | Agriculture, Developed | | X | | | 0.477 | 17.5 | 308.65 | 617.85 | X | |
| | | | | | | Agriculture, Developed | | X | | | 0.532 | 14.2 | 309.68 | 621.83 | | |

| | | | | | | | | | | | | | | | |
|---|---|---|---|---|---|---|---|---|---|---|---|---|---|---|---|
| *Hippodamia convergens* | X | | | | | | | | | | 0.269 | 9.86 | 462.17 | 922.1 | |
| | X | X | | | | | | | | | 0.514 | 32 | 450.46 | 885.6 | |
| | | | X | X | X | Agriculture, Forest, Developed | X | | | | 0.625 | 33.6 | 424.46 | 838.246 | |
| | | | X | X | X | Agriculture, Forest, Developed | | X | | | 0.632 | 34.8 | 420.75 | 834.47 | |
| | | | X | X | X | Agriculture, Forest, Developed | | X | X | X | 0.199 | 36.8 | 417.5 | 829.69 | |
| | | | X | X | X | Agriculture, Developed | | X | X | X | 0.247 | 35.7 | 419.17 | 830.65 | |
| | | | X | X | X | Developed | | X | X | X | 0.21 | 35.4 | 418.83 | 828.24 | |
| | | | X | | X | Developed | | X | X | X | 0.303 | 33.7 | 418.28 | 829.32 | |
| | | | X | | X | Developed | | | X | X | 0.463 | 32.8 | 420.09 | 829.56 | X |
| | | | X | | | Developed | | | X | X | 0.322 | 30 | 421.74 | 835.88 | |
| *Chilocorus stigma* | X | | | | | | | | | | 0.0185 | 28.6 | 248.46 | 497.3 | |
| | X | X | | | | | | | | | 0.0261 | 30.1 | 242.65 | 510.23 | |
| | | | X | X | X | Agriculture, Forest, Developed | X | | | | 0.22 | 27.6 | 244.11 | 494.52 | |
| | | | X | X | X | Agriculture, Forest, Developed | | X | | | 0.31 | 29.1 | 241.33 | 491.33 | |
| | | | X | X | X | Agriculture, Forest, Developed | | X | X | X | 0.3 | 29.1 | 243.02 | 495.51 | |
| | | | | X | X | Agriculture, Forest, Developed | | X | | | 0.312 | 28.8 | 240.78 | 489.01 | |
| | | | | X | X | Forest, Developed | | X | | | 0.326 | 28.6 | 242.71 | 486.52 | |
| | | | | X | X | Forest | | X | | | 0.271 | 27.1 | 242.45 | 486.54 | X |
| | | | | | X | Forest | | X | | | 0.251 | 24.6 | 243.49 | 488.57 | |

**Figure S3.1.** Spatial distribution of *Adalia bipunctata* in the state of Ohio, adjusted for sampling effort, as predicted by a negative binomial Generalized Additive model. Isolines indicate areas of similarity between predicted captures. A) All data, 1930- present; B) 1930-1940; C) 1950-1970; D) 1980-2000; E) 2010-2018.

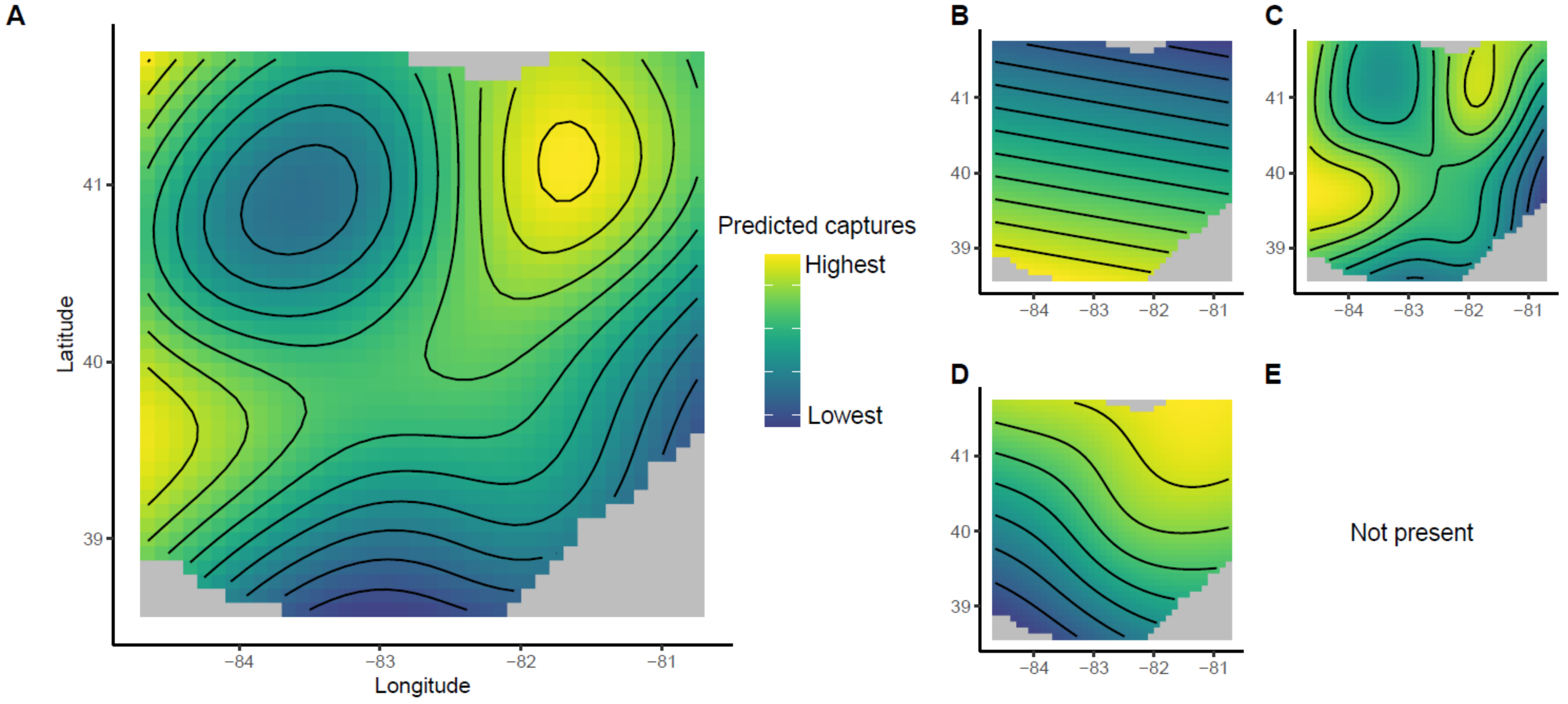

**Figure S3.2.** Spatial distribution of *Coccinella novemnotata* in the state of Ohio, adjusted for sampling effort, as predicted by a negative binomial Generalized Additive model. Isolines indicate areas of similarity between predicted captures. A) All data, 1930-present; B) 1930-1940; C) 1950-1970; D) 1980-2000; E) 2010-2018.

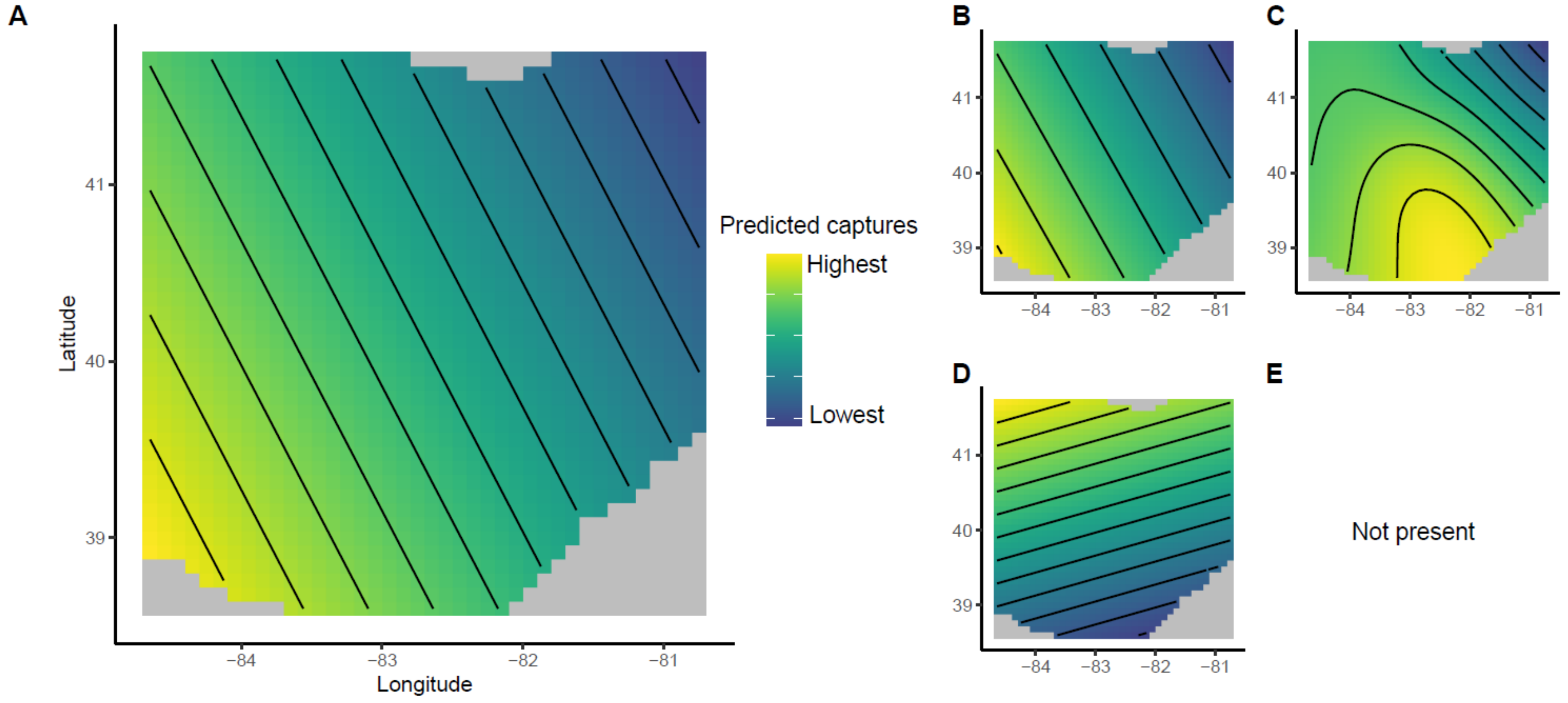

**Figure S3.3.** Spatial distribution of *Coleomegilla maculata* in the state of Ohio, adjusted for sampling effort, as predicted by a negative binomial Generalized Additive model. Isolines indicate areas of similarity between predicted captures. A) All data, 1930-present; B) 1930-1940; C) 1950-1970; D) 1980-2000; E) 2010-2018.

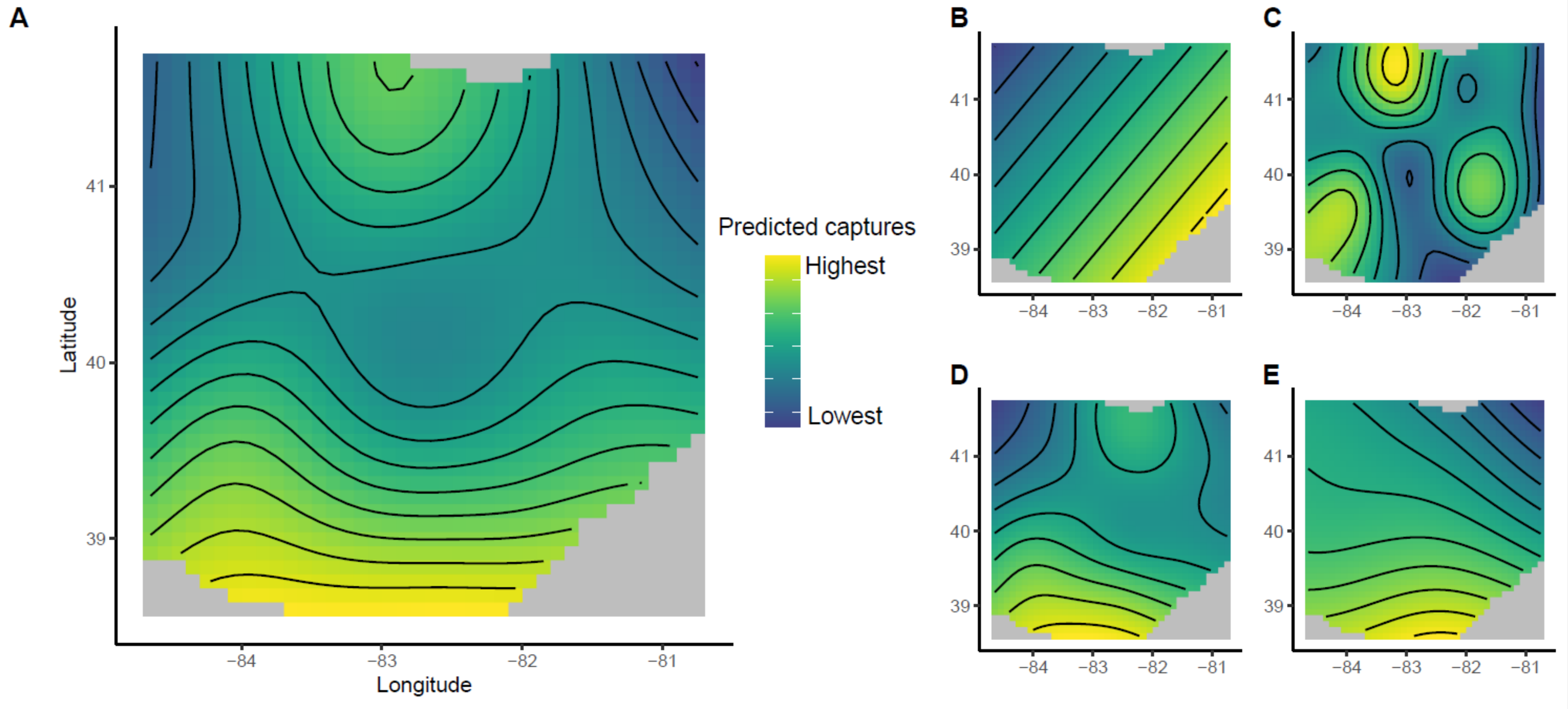

**Figure S3.4.** Spatial distribution of *Hippodamia convergens* in the state of Ohio, adjusted for sampling effort, as predicted by a negative binomial Generalized Additive model. Isolines indicate areas of similarity between predicted captures. A) All data, 1930-present; B) 1930-1940; C) 1950-1970; D) 1980-2000; E) 2010-2018.

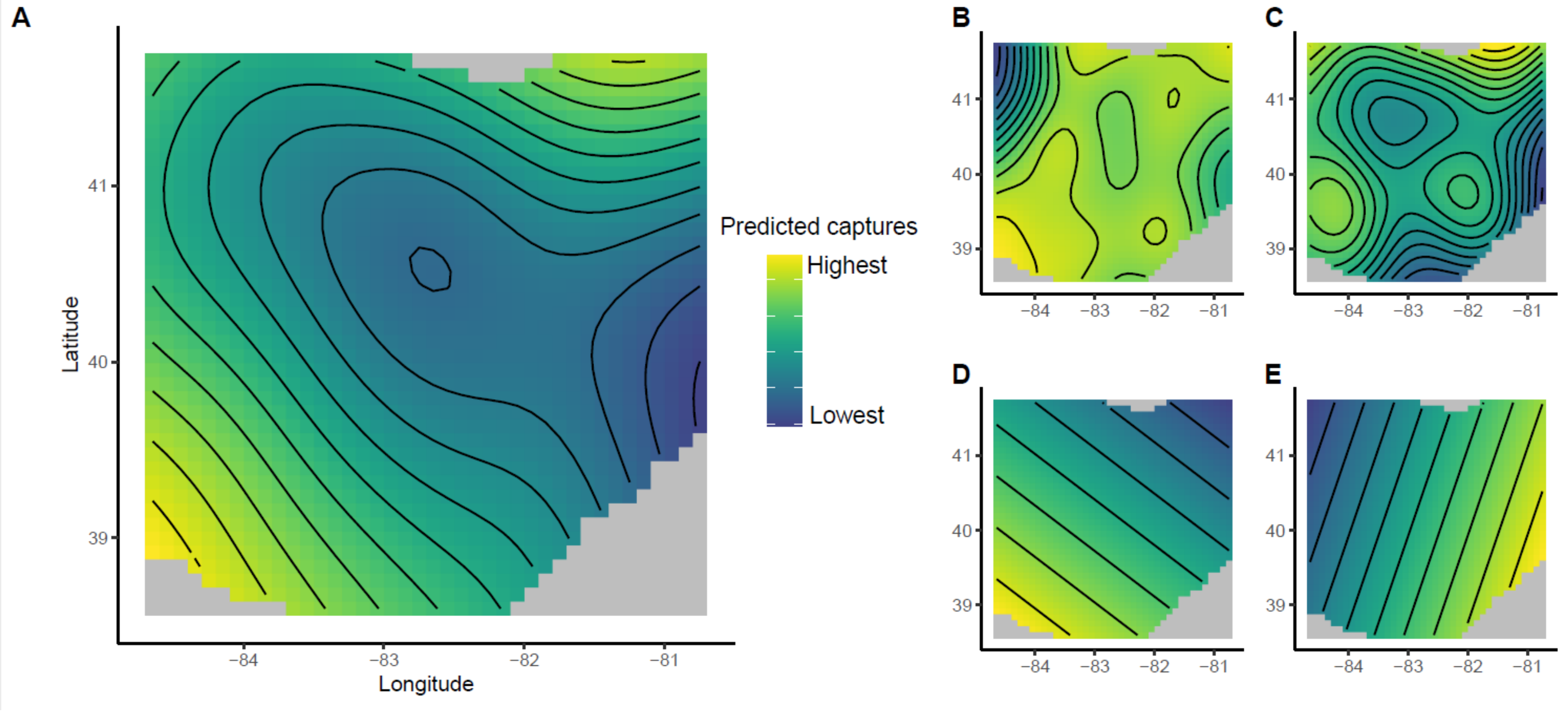

**Figure S3.5.** Spatial distribution of *Chilocorus stigma* in the state of Ohio, adjusted for sampling effort, as predicted by a negative binomial Generalized Additive model. Isolines indicate areas of similarity between predicted captures. A) All data, 1930- present; B) 1930-1940; C) 1950-1970; D) 1980-2000; E) 2010-2018.

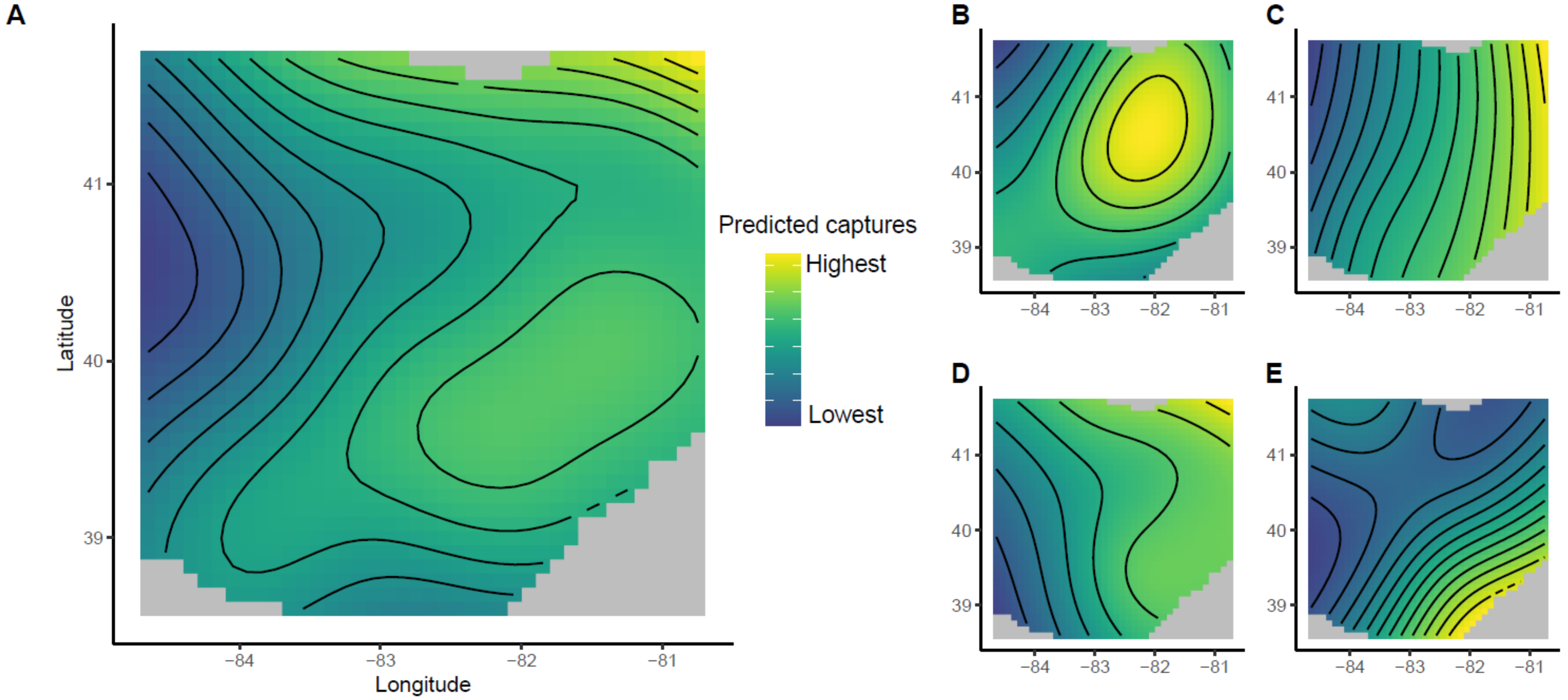
